# RNA degradation by DIS3 is a necessary step in the resolution of backtracked transcription complexes

**DOI:** 10.64898/2026.02.02.703186

**Authors:** Elin Enervald, Marcel Tarbier, Shruti Jain, Indranil Sinha, Akshit Jain, Jordi Planells, Toni McHugh, Neus Visa

**Affiliations:** Department of Molecular Biosciences, Wenner-Gren Institute, Stockholm University, SE-106 91 Stockholm, Sweden; Science for Life Laboratory, Department of Immunology, Genetics and Pathology, Rudbecklaboratoriet, Uppsala University, SE-751 85 Uppsala, Sweden; Department of Laboratory Medicine, Karolinska Institute, SE-171 77 Stockholm, Sweden; Instituto de Biotecnología y Biomedicina (BiotecMed), Departamento de Biología Celular, Universidad de Valencia, ES-46100, Spain; Light Microscopy Core, Discovery Research Platform for Hidden Cell Biology, University of Edinburgh, EH9 3FE, Scotland, UK

**Keywords:** RNA exosome, RNA polymerase II, transcription elongation, backtracking, UV-induced DNA damage

## Abstract

The RNA exosome is known to participate in transcription, but the contribution of its ribonuclease activities to this process remains unclear. Here we investigated the role of DIS3, one of the exosome ribonucleases, in transcription by RNA polymerase II (RNAPII). Rapid depletion of DIS3 reduced RNA synthesis and induced RNAPII elongation defects that were exacerbated by UV irradiation, a treatment that generates transcription-blocking DNA lesions and promotes RNAPII backtracking. Notably, DIS3 itself was redistributed following UV irradiation in a manner that closely paralleled RNAPII dynamics, which suggested that DIS3 acts in concert with the transcription machinery. We also investigated whether RNA degradation by DIS3 was required for transcription elongation and found that the 3’–5’ exoribonucleolytic activity of DIS3, but not its endonucleolytic activity, is essential for efficient transcription elongation. More specifically, DIS3 degrades the 3’ ends of backtracked RNA, as shown by sequencing of RNA fragments released by TFIIS-induced transcript cleavage *in vitro*. This DIS3-dependent degradation of backtracked RNA is critical for resolving stalled RNAPII complexes and enabling productive transcription elongation.

**Graphical abstract:** 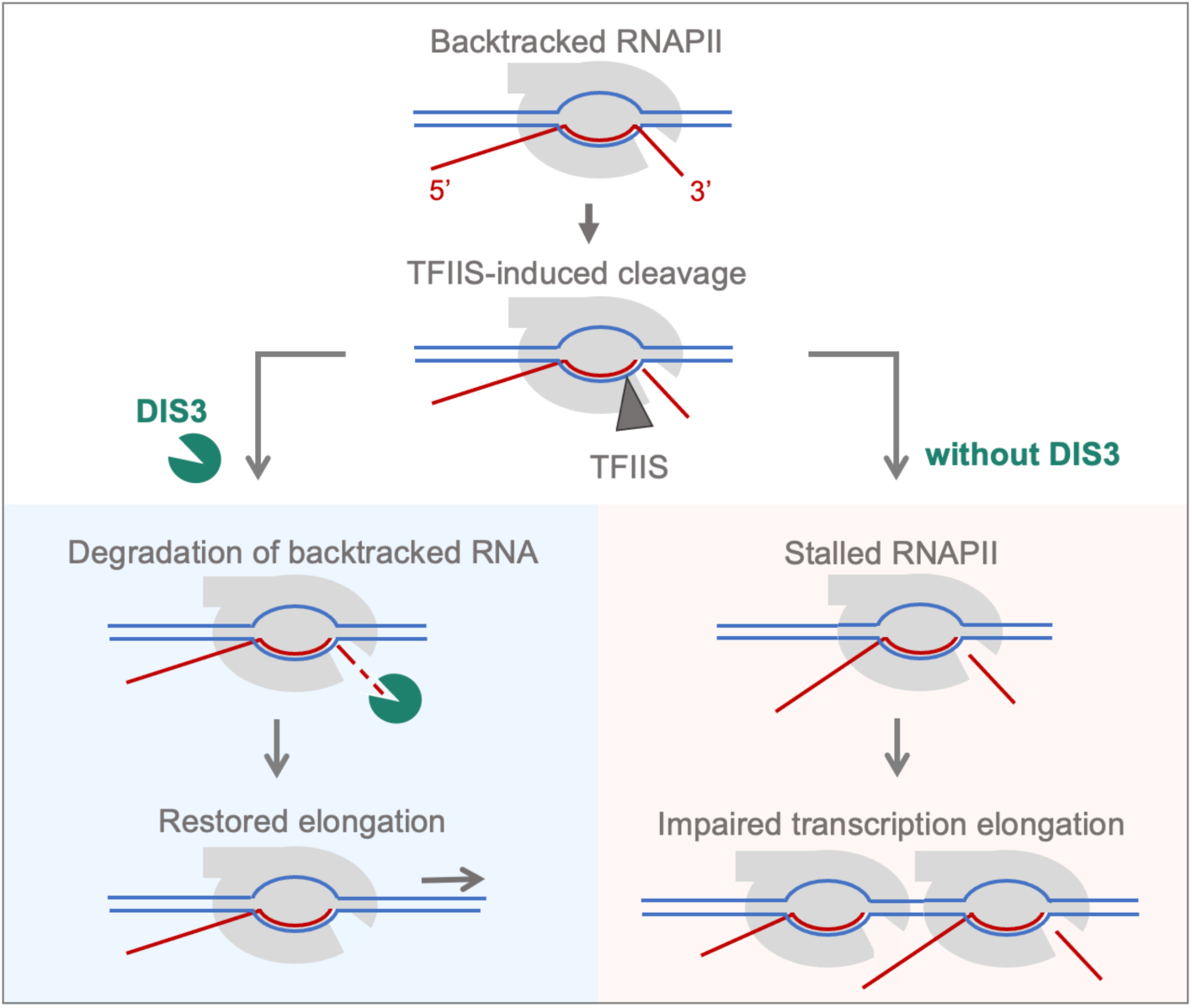

**Highlights:**

- DIS3 depletion leads to severely reduced rates of RNA synthesis in human cells.
- Transcription-blocking lesions reveal a critical role for DIS3 in RNA polymerase II elongation.
- The 3’ to 5’ exoribonuclease activity of DIS3 is necessary for transcription elongation.
- DIS3-mediated degradation of backtracked RNA is required for the resolution of stalled transcription complexes.

## Introduction

The RNA exosome is one of the major RNA degradation systems in eukaryotic cells. In both in archaea and eukaryotes, the RNA exosome is composed of a core made of nine subunits that form a barrel-shaped structure. In eukaryotes, the core is catalytically inactive and associates with two hydrolytic ribonucleases: EXOSC10 and DIS3 (Tomecki et al. 2010, Chlebowski et al. 2013, Januszyk and Lima 2014). EXOSC10 is a 3’-5’ exoribonuclease. DIS3 has a PIN domain that confers endoribonucleolytic activity and a RNB domain with 3’-5’ exoribonucleolytic activity (Lebreton et al. 2008, Schaeffer et al. 2009). In the cell nucleus, the RNA exosome contributes to the processing of many non-coding RNAs (ncRNAs), including rRNAs and snoRNAs, plays a major role in the quality control of mRNA biogenesis, and is responsible for the degradation of cryptic and pervasive transcripts (reviewed by Schmid and Jensen 2008, Jensen et al. 2013, Chlebowski et al. 2013, Garland and Jensen 2024). Due to its action on a plethora of different RNA substrates, the RNA exosome is essential for a diversity of biological processes, including gene regulation, embryonic development, DNA repair and cell senescence (Lloret-Llinares et al. 2018; Petit et al. 2022; Domingo-Prim et al. 2019, Han et al. 2024).

The exosome catalytic subunit DIS3 is encoded by one of the most recurrently mutated genes in multiple myeloma (Manier et al. 2017, Walker et al. 2018) and recent studies have shown that DIS3 is required for the maintenance of genome integrity (Gritti et al. 2022, Mérida-Cerro et al. 2025, Kuliński et al. 2025). However, depletion of DIS3 *per se* does not affect the overall efficiency of DNA double-strand break repair (Domingo-Prim et al. 2019) and the mechanisms underlying the genome instability observed in cells with altered DIS3 levels are not fully understood.

The functions of the RNA exosome have been thoroughly investigated in the context of RNA processing, quality control of gene expression, and sorting of spurious transcripts (reviewed by Houseley et al. 2006, Garland and Jensen 2024). Extensive experimental evidence also supports a role for the RNA exosome in transcription by RNA polymerase II (RNAPII). Transcriptomic analyses of mouse embryonic stem cells showed that EXOSC3 and EXOSC10 are needed for the resolution of harmful transcription-coupled secondary DNA structures (Pefanis et al. 2015). Subsequent studies in the EXOSC3-deficient mouse cells suggested that the RNA exosome cooperates with the nuclear exosome targeting (NEXT) complex and the Integrator (INT) to promote promoter-proximal termination of RNAPII (Torre et al. 2023). In human cells, the RNA exosome forms stable complexes with the transcription factor MYCN, regulates the expression of the MYCN-dependent genes, and maintains productive transcription elongation (Papadopoulos et al. 2022, Papadopoulos et al. 2024). Moreover, through its association with the ARS2-ZC3H4-NEXT complex, the RNA exosome has been implicated in transcription termination (Rouvière et al. 2023) and linked to a DNA-dependent transcription termination mechanism that preferentially targets promoter-proximal transcripts of specific genes containing T-rich sequence elements (Davidson et al. 2024).

While it is evident that the RNA exosome is involved in transcription, the underlying molecular mechanisms and the extent to which its catalytic activities contribute to RNAPII-mediated transcription remain poorly understood. Early studies in *Schizosaccharomyces pombe* led to the proposal that DIS3 acts as a ’fail-safe’ mechanism for transcription termination by cotranscriptionally degrading RNA at backtracked and stalled RNAPII complexes, a process described as the *reverse torpedo model* (Lemay et al. 2014). RNA profiling of mammalian cells expressing a PIN-RNB double mutant of DIS3 also supported a role for DIS3 in degrading potentially premature RNAPII termination products (Szczepińska et al. 2015). However, the contribution of DIS3 to the resolution of backtracked complexes has remained controversial primarily because RNAPII backtracking was estimated to span only 2–14 base pairs, as inferred from the size of endonucleolytic fragments generated by TFIIS-stimulated cleavage (Izban and Luse, 1993). The 3′ end of a short cleavage product would remain within the RNAPII funnel (Wang et al. 2009) and would be inaccessible to nucleoplasmic ribonucleases.

The recent use of long-range cleavage sequencing (LORAX-seq), a method for sequencing backtracked RNAs with nucleotide resolution *in vivo*, has revealed that persistent backtracking events spanning 20-40 nt are not unusual in human cells (Yang et al. 2024). The occurrence of long range, persistent backtracking raises the possibility that the RNA exosome contributes to RNAPII transcription through exonucleolytic cleavage of backtracked RNA by DIS3. Here, we have revisited this hypothesis and analyzed the role of DIS3 in transcription. By using UV irradiation to produce transcription-blocking lesions and in this way challenge the backtracking machinery, we demonstrate that rapid depletion of DIS3 results in RNAPII accumulation at transcription start sites (TSSs) and reduced elongation rates. Using LORAX-seq we show that DIS3 degrades backtracked RNA, and by analyzing catalytically inactive mutants of DIS3 we provide direct evidence that its exoribonucleolytic activity is required for efficient transcription elongation. Collectively, our present observations uncover a novel function for DIS3 in RNAPII transcription in human cells. Aberrant RNAPII stalling, resulting from impaired DIS3-mediated resolution of backtracked transcription complexes, may constitute a major source of genomic instability in DIS3-deficient cells.

## Results

### DIS3 is required for efficient transcription elongation

We carried out loss-of-function studies using an auxin-inducible degron cell system (DIS3-AID) developed by Davidson et al. (2019) to rapidly degrade DIS3 in human HCT116 cells (Suppl. Fig. 1A). In order to determine whether DIS3 is required for transcription, we conducted nascent RNA measurements using metabolic labeling with 5-EU and subsequent quantification of nascent transcripts by fluorescence microscopy in auxin-treated DIS3-AID cells and in control cells treated with DMSO. We recorded 5-EU labeling in randomly selected nuclear regions outside the nucleolus to avoid confounding effects from rRNA synthesis by RNAPI. Control cells showed sustained transcription over the course of the experiment while DIS3-depleted cells showed a progressive decrease in transcription throughout the course of the experiment (Fig. 1A). The transcriptional defect resulting from DIS3 depletion was rescued by overexpression of a doxycycline-inducible DIS3 (Suppl. Fig. S1B-D).

**Figure 1.**
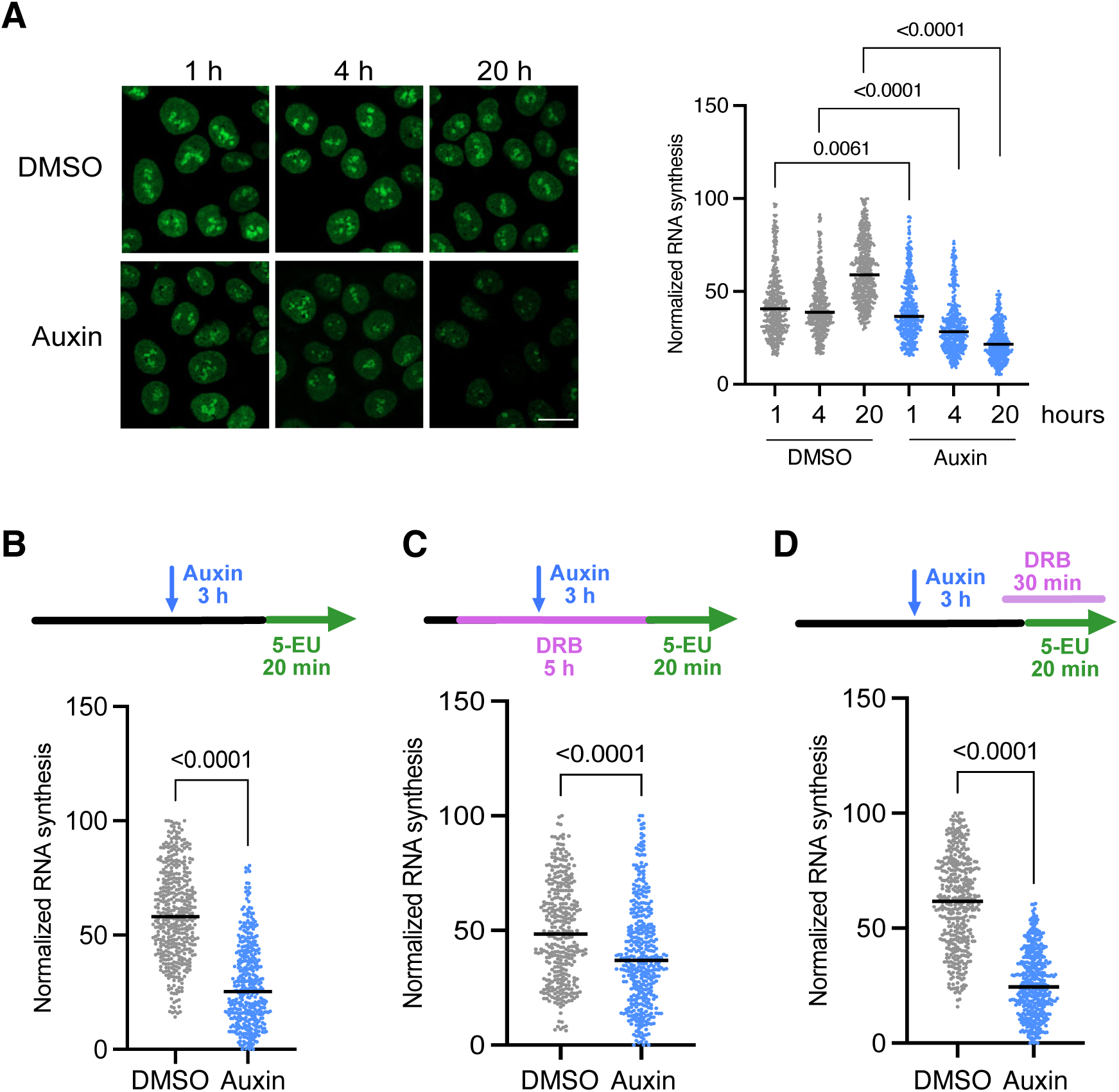
DIS3 is needed for efficient RNA synthesis. (A) DIS3-AID cells were treated either with DMSO (control) or auxin for rapid depletion of DIS3. Nascent RNA was quantified at the indicated time points by 5-EU incorporation (20 min pulse) and labeling. The figure shows representative images (left) and quantitative analysis (right). The absolute 5-EU signal was normalized within each experiment. Black lines indicate the median. The plot shows 5-EU normalized signal for cells in the 5^th^ - 95^th^ percentile from three independent experiments. Number of analyzed cells in each condition: DMSO: 1h *n* = 459, 4h *n* = 445, 20h *n* = 477; Auxin: 1h *n* = 434, 4h *n* = 421, 20h *n* = 441. A Mann-Whitney test was used to test statistical significance. The scale bar represents approximately 10 μm. (B-D) Transcription analysis in the presence of DRB to uncouple transcription elongation from initiation. In all cases, RNA synthesis was quantified by 5-EU after a 20 min incorporation pulse as in (A). (B) shows a control experiment without DRB treatment to measure transcription in DIS3-AID cells, as in (A). In (C), DIS3-AID cells were treated with DRB for 5 h, DRB was washed out, and the cells were treated with 5-EU for pulse-labeling in fresh medium. In (D), DRB was added 10 min prior to addition of 5-EU for pulse-labeling. Normalized transcription levels were determined as in A. The plots show compiled data from three independent experiments. Number of analyzed cells in each condition: *No treatment* DMSO: *n* = 429, Auxin: *n* = 401; *DRB block and wash out* DMSO: *n* = 348, Auxin: *n* = 396; *DRB block* DMSO: *n* = 416, Auxin: *n* = 453. A Mann-Whitney test was used to test statistical significance.

To identify which step in the RNAPII transcription cycle was affected by DIS3 depletion, we used the CDK9-inhibitor 5,6-dichloro-1-b-D-ribofuranosylbenzimidazole (DRB) in combination with nascent RNA quantification by 5-EU incorporation. The cells were first cultured for 3 hours in the presence of either DMSO or auxin to deplete DIS3. Following this incubation, the cells were subjected to one of three different treatments as illustrated in Fig. 1B-D. In control experiments without DRB, depletion of DIS3 significantly reduced nascent RNA synthesis (Fig. 1B). In DRB block and wash-out experiments (Fig. 1C), we investigated whether DIS3 was involved in the transition from transcription initiation to elongation. We first treated the cells with DRB for 5 hours, a treatment that allows already elongating RNAPII compelxes to complete their run-off while preventing new polymerases from engaging into elongation. Following this incubation, DRB was removed to allow the release of RNAPII complexes paused at the TSS. In this conditions, nascent transcription in DIS3-depleted cells was significantly lower than in DMSO-treated cells, which revealed that DIS3 is required for an efficient initiation-to-elongation transition (Fig. 1C).

The third treatment was a DRB block aimed to investigate the role of DIS3 in elongation. We treated the cells with DRB for 10 min prior to the 20-min pulse labeling with 5-EU. Also in this case, the depletion of DIS3 led to significantly reduced nascent RNA levels, indicating a transcription elongation defect (Fig. 1D). This defect was more severe than that observed in the DRB washout experiment, suggesting that the primary consequence of DIS3 depletion is a general reduction of RNAPII elongation.

### Transcription-blocking lesions reveal a critical role for DIS3 in RNAPII elongation

Next, we generated high-throughput RNAPII profiles by CUT&Tag to further characterize the effects of DIS3 depletion on transcription. We used antibodies against the elongating RNAPII phosphorylated at the Ser2 of the RPB1 C-terminal domain (RNAPII S2) and against the initiating/paused RNAPII phosphorylated at Ser5 (RNAPII S5). Metagene plots of normalized coverage for RNAPII S2 and S5 showed that depletion of DIS3 resulted in increased RNAPII levels near the transcription start site (TSS) and in the promoter-proximal part of the gene body (Fig. 2A). Conversely, DIS3-depleted cells showed lower S2 and S5 RNAPII levels in the distal part of the gene body and downstream of the cleavage site (transcription end site, TES), which is consistent with an elongation defect that impairs RNAPII processivity and limits the number of transcription complexes that reach the end of the gene.

**Figure 2.**
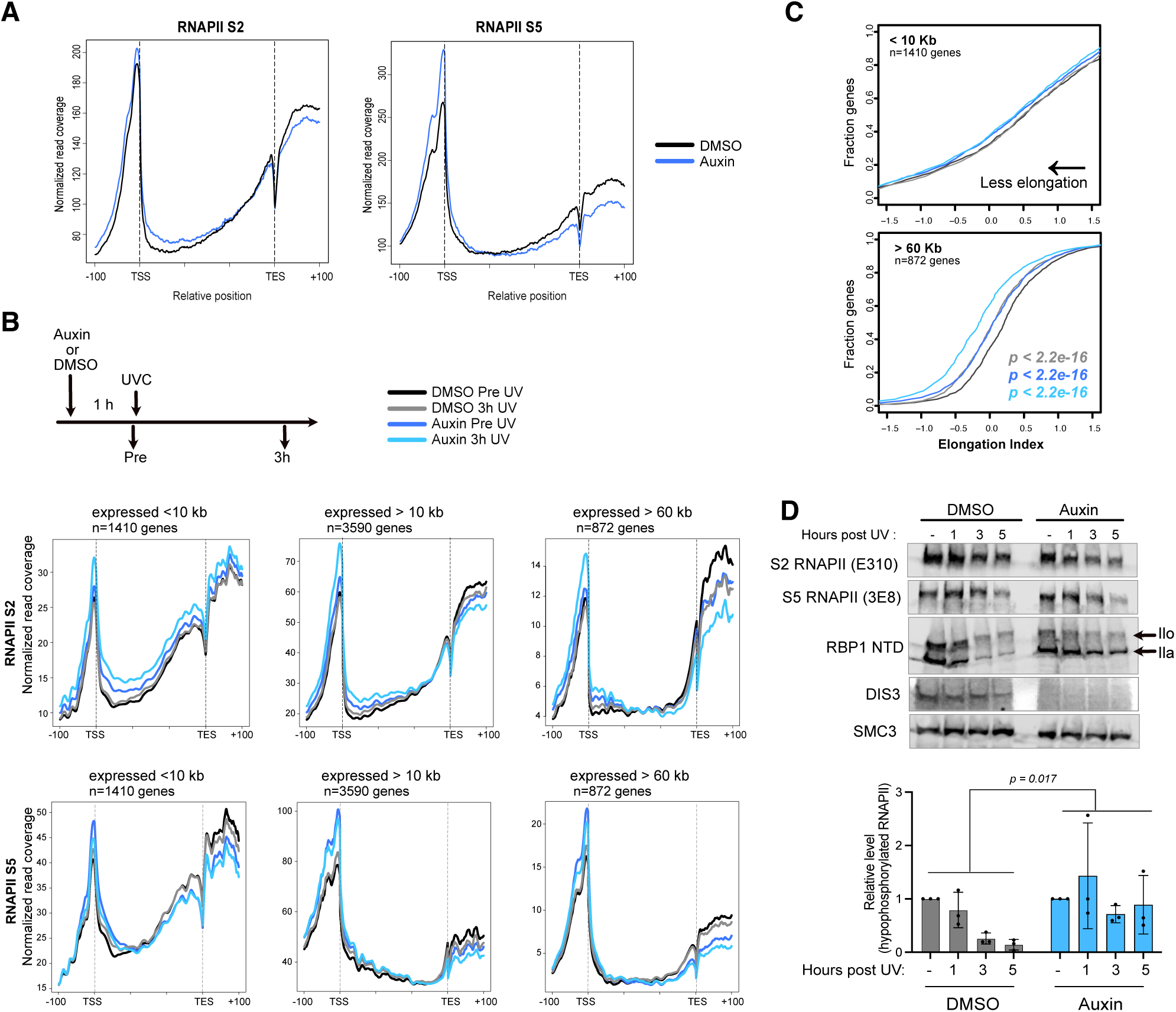
DIS3 depletion inhibits RNAPII elongation. (A) DIS3-AID cells were treated with either DMSO (control) or auxin for rapid depletion of DIS3. CUT&Tag was carried out after 4 hours of auxin treatment to analyze the effect of DIS3 depletion on RNAPII elongation using antibodies against RNAPII S2 and S5, as indicated. The plots show normalized read coverage. n=5002 genes. (B) DIS3-AID cells were treated with either DMSO (control) or auxin for 1h prior to UVC irradiation. Irradiation dose was 8 J/m^2^ UVC. Metagene plots show the CUT&Tag normalized read coverage for RNAPII S2 and S5 over gene bodies ±100 bp in control cells (DMSO) or DIS3-depleted cells (Auxin) with or without irradiation, as indicated. Only expressed genes (>10 fpkm) according to data from Davidson et al. (2019) are included in the analysis. Genes of different lengths are plotted separately. (C) Elongation indexes (EI) for RNAPII S2 in the same four experimental conditions analyzed in B. EI was defined as the ratio of normalized RNAPII S2 coverage in the distal half of the genes over the normalized RNAPII S2 coverage in the promoter-proximal half of the gene. P-values for each condition compared to the control sample (DMSO, pre-UV) result from a one-sided Mann-Whitney U test / Wilcoxon rank-sum test. (D) Western blot analysis of chromatin-bound proteins extracted from DIS3-AID cells treated with either DMSO or auxin prior to irradiation with 20 J/m^2^ UVC. SMC3 was used as a loading control for normalization purposes. Quantification from three independent experiments is plotted in the bottom part of the figure. A two-way Anova was used for statistical testing.

Based on the transcriptional elongation defects described above and on the changed RNAPII occupancy patterns caused by DIS3 depletion, we hypothesized that DIS3 might be required for the resolution of stalled transcription complexes. If this is the case, alterations in RNAPII occupancy in DIS3-depleted cells should be exacerbated by treatments that induce RNAPII backtracking and stalling. In order to test this possibility, we irradiated DIS3-AID cells with UV light to produce transcription blocking lesions in the DNA. RNAPII complexes stalled at DNA lesions are known to be either backtracked, disassembled, or degraded via ubiquitylation and proteasomal degradation (reviewed by Nieto Moreno et al. 2023).

We analyzed the normalized coverage for RNAPII S2 and S5 in control cells and in DIS3-depleted cells, before UV irradiation and three hours after irradiation (Fig. 2B). In short genes (<10 kb), UV irradiation of DIS3-depleted cells resulted in increased RNAPII S2 levels throughout the gene but reduced RNAPII S5 in downstream sequences. In longer genes (>10 kb and > 60 kb), DIS3 depletion resulted in increased RNAPII levels in the promoter proximal part of the genes but reduced occupancy in the 3’ regions and downstream of the TES. For a more quantitative description of the effect of DIS3 in transcription elongation, we calculated the Elongation Index (EI) of RNAPII S2 (see Methods). A decreased EI indicates that fewer RNAPII molecules reach the distal part of the gene body. In control cells (DMSO-treated), the EI was significantly lower 3 hours post UV than before irradiation, as expected. Auxin-treated cells showed even lower EIs, both in irradiated and non-irradiated cells, compared to the corresponding controls treated with DMSO (Fig. 2C). Not surprisingly, elongation defects were most pronounced in long genes (Fig. 2B-C).

ChIP-qPCR experiments in U2OS cells confirmed transcription elongation defects associated with DIS3 depletion (Suppl. Fig. S2A).

In another series of experiments, we analyzed the changes in RNAPII bound to chromatin by Western blot analysis of chromatin fractions using antibodies against the N-terminal domain of RPB1 (NTD). The global levels of engaged RNAPII gradually decrease following UV damage, and this effect was also observed in DIS3-depleted cells (Fig. 2D). The pre-initiating hypophosphorylated RNAPIIa was also reduced in DMSO-treated cells (control) following UV irradiation, correlating with the UV-induced clearance of RNAPII from promoter regions described in earlier reports (Steurer et al. 2022). However, in DIS3-depleted cells, a larger fraction of hypophosphorylated RNAPIIa remained bound to chromatin after UV irradiation (Fig. 2D).

We also asked whether RNAPII retention on chromatin was a general response to DIS3 depletion. Western blot analyses of RNAPII in chromatin fractions of U2OS and Hela cells showed increased RNAPII levels after DIS3 depletion by RNA interference (Suppl. Fig. S2B). We concluded that RNAPII retention on chromatin represent a general response to DIS3 depletion, not unique to the DIS3-AID cell line.

In summary, the analysis of RNAPII occupancy confirmed that DIS3 is necessary for proper transcription elongation, both under normal conditions and when the transcription machinery is challenged by transcription blocking lesions.

### UV-induced redistribution of DIS3 recapitulates RNAPII dynamics

Previous studies have shown that the loss of DIS3 results in accumulation of thousands of transcripts (Davisdon et al. 2019) and affects DNA:RNA hybrid levels in both mammalian and yeast cells (Gritti et al. 2022, Mérida-Cerro et al. 2025). Thus, we hypothesized that unresolved DNA:RNA hybrids could be the underlying cause of the RNAPII elongations defects reported above. We carried out slot blot experiments to quantify DNA:RNA hybrid levels in the DIS3-AID cells (Suppl. Fig. S3A). We also quantified DNA:RNA hybrids in a U2OS cell line expressing the hybrid-binding domain of RNase H fused to GFP (Suppl. Fig. S3B). None of the experiments provided support for this hypothesis.

In order to get mechanistic insight into the role of DIS3 in transcription, we asked whether the association of DIS3 with transcription units was affected by UV irradiation. It is well established that exposure to UV light alters the distribution of RNAPII (Lavigne et al. 2017, Van den Heuvel et al. 2021, reviewed by Nieto Moreno et al. 2023), but the impact of UV irradiation on the genomic distribution of DIS3 has not been investigated. We generated ChIP-seq profiles of DIS3 in U2OS cells that express a HA-tagged DIS3 protein (DIS3^FLAGHA^). We also analyzed the total occupancy of RNAPII using an antibody against the N-terminal domain of RPB1 (NTD) (Fig. 3 and Suppl. Fig. S4). Average RNAPII occupancy at transcription start sites (TSSs) was significantly decreased 3 hours after UV irradiation, consistent with previous reports, and was notably recovered 8 hours after irradiation (Fig. 3A-B). A more detailed analysis of RNAPII redistribution by K-means clustering revealed a considerable diversity in the transcriptional response to UV irradiation (Fig. 3C). The genes in cluster 2 (n=1054 genes) showed a significant reduction of RNAPII at the transcription start site (TSS) following irradiation (Fig. 3C), with a remarkable recovery at 8 hours (Fig. 3D). Notably, these genes displayed higher levels of both RNAPII and DIS3 in control cells than those in clusters 1 and 3 (Suppl. Fig. S4), indicating that they are highly expressed.

**Figure 3.**
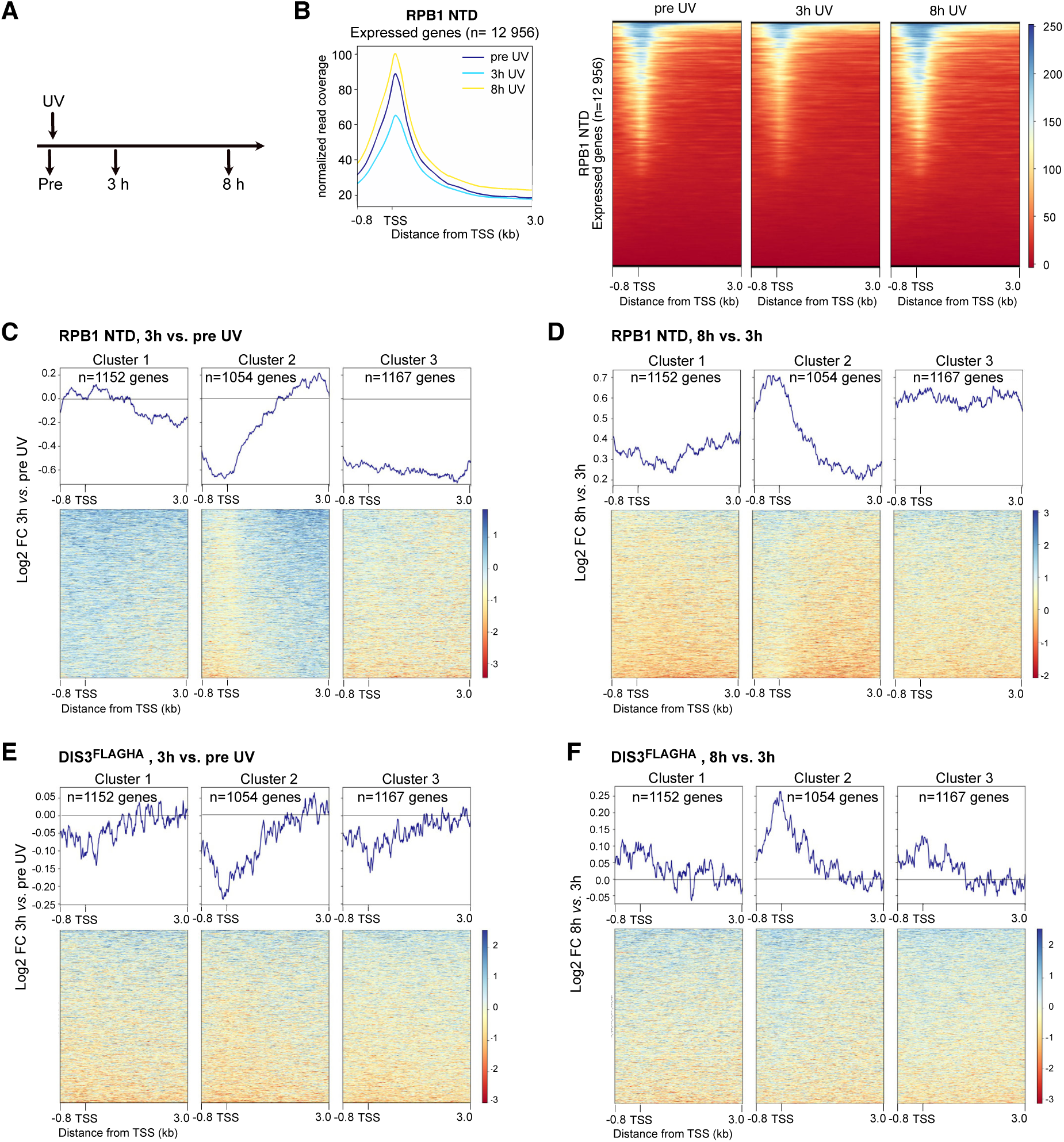
Redistribution of RNAPII and DIS3 in response to UV irradiation analyzed by ChIP-seq. A. Schematic of the UV irradiation experiment. B. ChIP-seq analysis of RNAPII in U2OS cells that express a HA-tagged DIS3 protein (U2OS-DIS3^FLAGHA^-WT) prior to irradiation (pre) and at different timepoints after UV irradiation (8 J/m^2^), as indicated, using an antibody against the RPB1 NTD. The image shows RPB1 NTD ChIP signal distribution across different UV-treatment timepoints, at the TSS and 3 kb into the gene body. The signal is expressed as average read coverage normalized to input. The corresponding heat maps are shown on the right-hand side of the image. Non-expressed genes were excluded from the analysis. C. K-means clustering analysis of RPB1 NTD fold change at 3 h after UV irradiation compared to non-irradiated cells. The plots include merged data from two replicates. The signal has been centered around the TSS as in (B). D. Metagene and heat map showing changes in RPB1 NTD occupancy at 8 h compared to 3h for the same clusters defined in (C). The plots show merged data from two replicates. (E, F) Metagenes and heat maps showing changes in DIS3 occupancy in the same clusters defined in (C). (E) shows DIS3 occupancy change at 3 h compared to pre-UV, and (F) shows change at 8 h compared to 3h. The plots show merged data from two replicates.

In non-irradiated cells, DIS3 was more abundant at transcription start sites (TSSs) than in gene bodies and showed an overall distribution similar to that of RNAPII (compare Suppl. Fig. S4A and S4B). Upon UV irradiation, DIS3 exhibited a redistribution that closely aligned with the changes observed in RNAPII (Fig. 3E-F). This similarity, which was particularly pronounced in highly expressed genes (cluster 2), suggests that DIS3 is associated with the transcription machinery and can act locally at the transcription site during elongation.

### The exoribonucleolytic activity of DIS3 is necessary for the degradation of backtracked RNA

The findings reported above suggest that DIS3 acts in concert with the transcription machinery and is required for efficient transcription elongation. Mechanistically, we hypothesized that DIS3 degrades backtracked RNA and that this degradation is required for the resolution of stalled transcription complexes. In order to test this hypothesis, we carried out LORAX-seq experiments (Fig. 4A) to isolate transcription complexes from DIS3-AID cells and sequence backtracked RNA products generated *in vitro* by TFIIS-induced cleavage (Suppl. Fig. S5). Backtracked RNAs isolated from DIS3-depleted cells were longer than those isolated from control cells. The average lengths for unique backtracked RNAs from auxin-treated and DMSO-treated cells were 30.9 and 27.8 nt, respectively (Fig. 4B and Suppl. Table S3), which demonstrates that DIS3 acts on backtracked RNAs.

**Figure 4.**
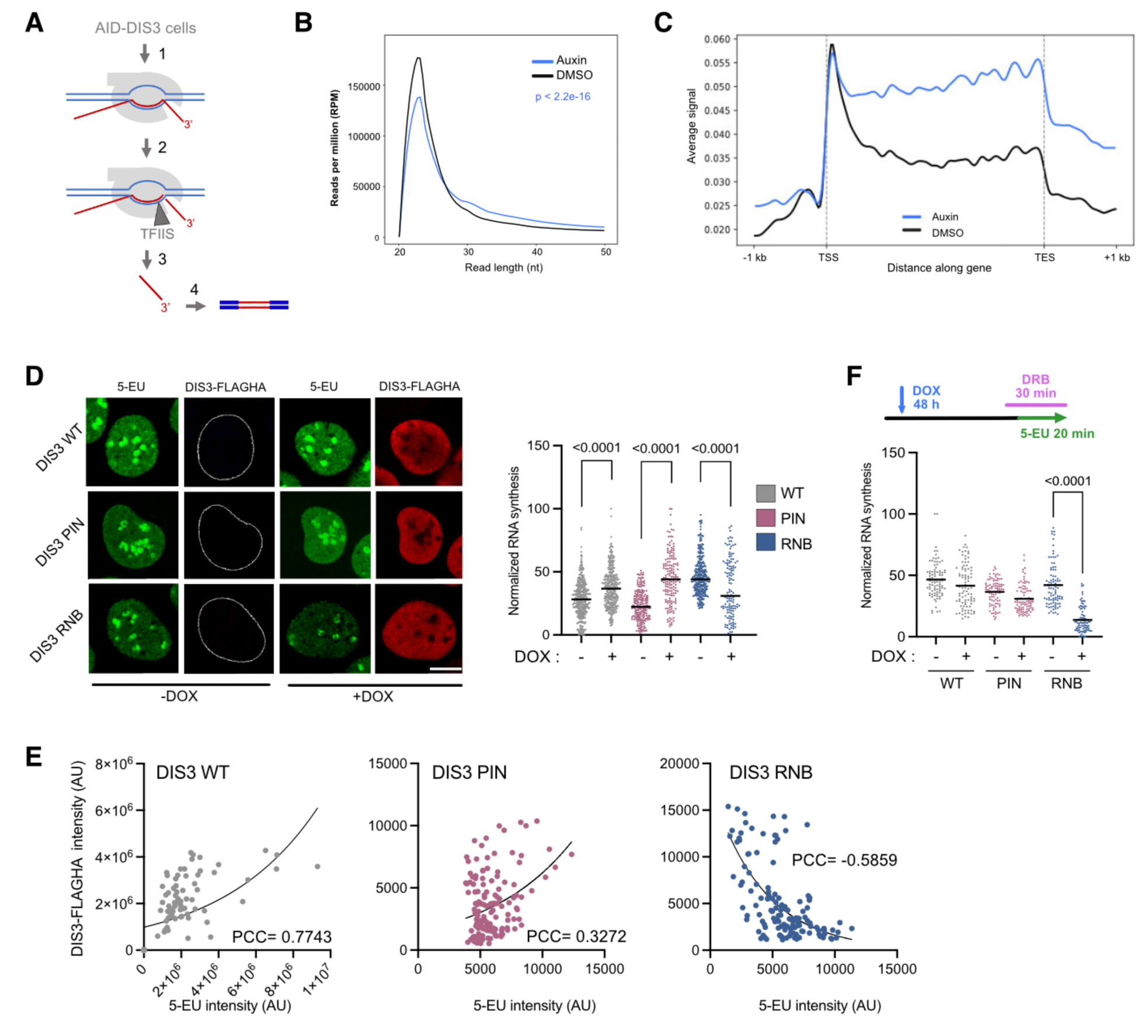
The exoribonuclease activity of DIS3 is needed for efficient transcription elongation and degradation of backtracked RNA. (A) Schematic of LORAX-seq. RNAPII complexes are immunoprecipitated from native chromatin preparations (1), the isolated chromatin is first treated with alkaline phosphatase to dephosphorylate the 5’ end of the nascent RNA, and then with TFIIS to release backtracked RNAs (2), the backtracked RNAs are purified (3) and sequenced (4). (B) Read length of backtracked RNAs analyzed by LORAX-seq. The plot shows read density of spike-in normalized RPMs (reads per million) in DMSO and Auxin treated cells. Only unique mapping reads are plotted. Data from two replicates per treatment were combined to generate merged distributions. Statistical comparisons between treatments were performed using Kolmogorov–Smirnov tests, with Benjamini–Hochberg correction for multiple testing. (C) Metagene plot showing the distribution of LORAX-seq signal from DMSO and Auxin treated cells across the gene bodies for expressed genes (>10 fpkm) with length <10kb. n=12228 transcripts, mapping to 1410 genes. (D) Nascent RNA was quantified by 5-EU incorporation in U2OS FLP-In T-rex cell lines overexpressing wild-type (WT) or mutant DIS3^FLAGHA^ variants. 5-EU labeling was carried out and quantified as in Fig. 1. Cells were counterstained with anti-HA antibody to quantify DIS3^FLAGHA^ expression. The figure shows representative images for each condition (top). The scale bars represent approximately 10 μm. The plot (bottom) includes the 5^th^-95^th^ percentile of all cells from three (DIS3 WT, DIS3 RNB) or two (DIS3 PIN) independent experiments. Number of cells analyzed in each condition: DIS3 WT: -DOX *n=*377, +DOX *n=*329; DIS3 PIN: -DOX *n=*292, +DOX *n=*193; DIS3 RNB: -DOX *n=*325, +DOX *n=*128. A Mann-Whitney test was used to test statistical significance. (E) Correlation analysis between levels of transcription (5-EU intensity) and DIS3- expression (HA) for each of the recombinant DIS3^FLAGHA^ variants, as indicated. Pearson’s correlation coefficients (PCC) are indicated. (F) Analysis of transcription elongation in the presence of DRB to quantify RNAPII elongation in cells overexpressing DIS3^FLAGHA^ variants. Quantification of 5-EU signal was carried out as in (D). The plot includes data from two independent experiments, *n*=88 cells analyzed in each condition. A Mann-Whitney test was used to test statistical significance. Significant *P* values are indicated in the figure.

We generated metagenes of normalized LORAX-seq signal in the proximity of the TSS. In agreement with previous studies (Yang et al. 2024), we observed a strong enrichment of backtracked RNA within 250 nt of the TSS, which presumably correspond to promoter-proximal pausing (Fig. 4C). Interestingly, in DIS3-depleted cells, we also observed significantly higher LORAX-seq signals in the gene bodies (Fig. 4C and Suppl. Fig. S5B). This finding futher supports the conclusion that DIS3 is required for the resolution of backtracked RNAPII.

The results reported above are consistent with the idea that the degradation of backtracked RNA by DIS3 is required for the resolution of backtracked RNAPII. For DIS3 to degrade backtracked RNA, the protein must be positioned near the RNAPII funnel domain, a surface of RNAPII that interacts with TFIIS (Kettenberger et al. 2004, Cheung and Cramer 2011). Proximity ligation assays (PLAs) revealed a strong interaction between DIS3 and TFIIS (Suppl. Fig. S6), which provides further support to the proposal that DIS3 acts on backtracked transcription complexes.

The function of DIS3 in degrading backtracked RNA would also require its 3’ to 5’ exoribonucleolytic activity. Thus we predicted that this activity is critical for effective RNAPII transcription. To test this prediction, we generated U2OS cells that expressed DIS3 variants under the control of a doxycyclin-inducible promoter and interrogated their contribution to transcription elongation. We induced expression of either wild-type DIS3 (referred to as DIS3 WT in Fig. 4D-F) or mutant versions carrying point mutations in the catalytic domains. One of the mutants, DIS3 D146N (DIS3 PIN), renders the endonucleolytic domain inactive. The other mutant, DIS3 D487N (DIS3 RNB), carries an inactive 3’-5’ exoribonuclease domain (Tomecki et al. 2014). Importantly, these mutations do not abrogate the nuclear localization of DIS3, and the distribution pattern of the mutant proteins in the cell nucleus resembles that of the endogenous DIS3 (Suppl. Fig. S7A). The exogenous DIS3 variants were overexpressed approximately 6-9 fold compared to the endogenous DIS3 protein (Suppl. Fig. S7B-C).

We induced the expression of the different doxocyclin-inducible DIS3 variants and analyzed 5-EU incorporation to assess the global level of RNAPII transcription in each condition. Overexpression of DIS3 WT or DIS3 PIN led to increased incorporation of 5-EU (Fig. 4D) and we observed a positive correlation between 5-EU incorporation and the expression levels of these DOX-inducible DIS3 variants (Fig. 4E). In contrast, overexpression of DIS3 RNB was negatively correlated with 5-EU incorporation. We concluded that DIS3 RNB has a dominant negative effect and inhibits nascent RNA synthesis. A DRB block experiment further confirmed the transcriptional defect of the DIS3 RNB mutant and directly linked transcription elongation with the 3’-5’ exoribonucleolytic activity of DIS3 (Fig. 4F).

Altogether, the results reported above show that degradation of backtracked RNAs by the RNB domain of DIS3 is a necessary step for efficient transcription elongation.

## Discussion

RNAPII backtracking is a well-documented phenomenon that often arises as a result of ribonucleotide misincorporation or when elongating transcription complexes run into roadblocks (Bondarenko et al. 2006, Churchman and Weissman 2011, James et al. 2017). Using a combination of loss of function methods, metabolic labeling, CUT&Tag analyses and sequencing of backtracked RNA products, we report that the ribonuclease DIS3 is essential for efficient transcription elongation in human cells. More precisely, our experiments show that the 3’-5’ exoribonucleolytic activity of DIS3 is required for the degradation of backtracked RNA and for the resolution of backtracked transcription complexes. This proposal is based on the following observations: 1) DIS3 depletion results in reduced RNA synthesis and reduced RNAPII elongation, 2) DIS3 and RNAPII show similar patterns of occupancy and redistribution upon UV irradiation, 3) the DIS3 RNB mutant reproduces the transcriptional defects observed in DIS3-depleted cells, 4) backtracked RNA fragments released after TFIIS-induced cleavage are longer in DIS3-depleted cells than in control cells, and 5) DIS3 depletion leads to the accumulation of backtracked transcription complexes in the chromatin.

Research from multiple laboratories has provided compelling evidence for the involvement of the RNA exosome in transcription elongation and termination in both yeast and mammalian cells (Lemay et al. 2014, Pefanis et al. 2015, Torre et al. 2023, Davidson et al. 2019, Papolopoulos et al. 2022, Davidson et al. 2024; Uhl et al. 2026). The present study shows that the function of the RNA exosome in transcription is linked to the 3′–5′ exoribonucleolytic activity of DIS3. Loss-of-function experiments targetting other exosome subunits have previously shown to affect transcription (Pefanis et al. 2015, Papadopoulos et al. 2022, Torre et al., 2023), which suggests that DIS3 requires the RNA exosome complex to enable productive elongation. Interestingly, EXOSC10 is also required for efficient transcription elongation, and depletion of TFIIS has been shown to increase the proximity between the RNAPII and EXOSC10 (Papadopoulos et al. 2022), which strengthens the functional involvement of the RNA exosome in RNAPII backtracking and suggests that the entire RNA exosome is involved in the resolution of backtracked transcription complexes.

We show that DIS3 occupancy in genic sequences parallels that of RNAPII, and that both proteins are redistributted in a similar manner in response to UV irradiation. These observations give support to a model in which DIS3 acts in close connection with the transcription machinery. Physical interactions between the RNA exosome, the nascent transcript, and elongating RNAPII have been previously reported (Andrulis et al. 2002, Hessle et al. 2012, Lubas et al. 2011, Papadopoulos et al. 2022). These interactions allow the exosome to accompany the transcription machinery as it progresses along the gene. Our present results, including PLA data, imply that these interactions place DIS3 near the RNAPII funnel to be able to access the 3’ end of backtracked transcripts.

We did not find evidence of substantial DNA:RNA hybrid accumulation in DIS3-depleted cells, suggesting that unresolved hybrids are unlikely to be the main cause of the observed transcription elongation defect. However, our experiments do not rule out the stabilization of relatively short hybrids formed between the 3’ end of the nascent RNA and the template DNA strand (Cheung and Cramer 2011). In experiments using the HBD-GFP, such hybrids would perhaps be concealed by the RNAPII complex and would thus be undetectable. The slot blot approach does not suffer from this limitation, and yet does not reveal any significant accumulation of DNA:RNA hybrids in DIS3-depleted cells. However, the slot blot quantifies the total amount of DNA:RNA hybrids in the genome. Any short hybrids that might form at backtracked RNAPII sites during transcription would constitute only a minor subset of the overall hybrid population, and fluctuations in their levels would likely remain below the detection limits of our assays.

The molecular interactions between the nascent RNA, the RNAPII pore, and the active site trigger loop, determine RNAPII pausing and backtracking (Palangat and Landick 2001, Cheung and Cramer 2011). Structural studies of backtracked RNAPII complexes have shown that the backtracked RNA interacts with a specific site within the RNAPII pore. This interaction is weak in cases of short backtracking, allowing RNAPII to resume elongation. In contrast, extensive backtracking causes the RNA to trap the trigger loop, which inhibits elongation. In this situation, TFIIS induces a conformational change that displaces the backtracked RNA from the backtracked site and induces the cleavage of the backtracked RNA (Kettenberger et al. 2004, Palangat et al. 2005, Cheung and Cramer 2011). DIS3 does not seem to be required for TFIIS-induced RNA cleavage, as similar amounts of eluted RNA products were recovered from DIS3-depleted and control cells in the LORAX-seq experiments. Thus, our results support a model in which DIS3 acts independently of TFIIS and engages in the degradation of the backtracked RNA, thereby promoting the resolution of backtracked RNAPII complexes. Further studies will be required to establish whether the degradation of backtracked RNA by DIS3 renders the RNAPII complex able to resume transcription or results in the termination of the stalled transcription complex, as proposed in *S. pombe* (Lemay et al. 2014).

DIS3 has been linked to genomic instability and cancer, with a particularly strong association to multiple myeloma (Walker et al. 2018, Chapman et al. 2011). In B cells, loss-of-function mutations of DIS3 drive oncogenic chromosomal translocations. Proposed mechanisms underlying DIS3-related mutagenesis include alterations in the pattern of CTCF and cohesin binding, and the establishment of abnormally open chromatin states that favor promiscuous editing by the activation-induced cytidine deaminase AID (Laffleur et al. 2021, Gritti et al. 2022, Kuliński et al. 2023). Interestingly, the genomic instability resulting from DIS3 loss is not restricted to B cells (Gritti et al. 2022, Mérida-Cerro et al. 2025), which indicates that DIS3 plays a more general role in safeguarding genome integrity. In this context, alterations in R-loop dynamics have been proposed as a source of genomic instability in cells with altered DIS3 levels (Gritti et al. 2022, Mérida-Cerro et al. 2025), which supports the idea that unresolved DNA:RNA hybrids constitute an AID-independent mechanism underlying the genomic instability linked to DIS3 loss. Our results point to another mechanism through which DIS3 depletion may drive genome instability: persistent RNAPII stalling in DIS3-depleted cells likely compromises genome integrity by increasing the frequency of transcription–replication conflicts.

## Resource availability

Requests for resources and reagents should be directed to the corresponding author, Neus Visa.

The ChIP-seq, CUT&Tag and LORAX-seq data produced in this study are available from NCBI Gene Expression Omnibus (GSE295184) and ArrayExpress (E-MTAB-16220, E-MTAB-16280).

The plugin used to count PLA foci is available at COIL-Edinburgh/Elin_Intensities_v2_1 (https://doi.org/10.5281/zenodo.17640956).

## Acknowledgements

We thank Steven West (Univ of Exeter, UK) and Daniel Durocher (University of Toronto, Canada) for providing DIS3-AID cells and U2OS Flp-In T-Rex cells, respectively. We thank Petra Beli (Gutenberg University, Mainz, Germany) for providing the pcDNA5/FRT/TO-GFP-HBD plasmid. We also thank BEA, the Bioinformatics and Expression Analysis core facility, which is supported by the board of research at the Karolinska Institute. The authors acknowledge support from the National Genomics Infrastructure (NGI) in Stockholm funded by Science for Life Laboratory, the Knut and Alice Wallenberg Foundation and the Swedish Research Council. We also thank Eduardo Sagredo and Vicent Pelechano (SciLifeLab, Stockholm) for access to a NextSeq 2000. Parts of the data handling were enabled by resources in project [NAISS 2024/22-1234] provided by the National Academic Infrastructure for Supercomputing in Sweden (NAISS) at UPPMAX and at the PDC Center for High Performance Computing, KTH. These resources are funded by the Swedish Research Council through grant agreement no. 2022-06725 and 2016-07213. We thank the Imaging Facility at Stockholm University (IFSU) for support with microscopy. This work was supported by grants from The Swedish Research Council [grants 2019-03853] and The Swedish Cancer Society [grant 19 0258 Pj] to NV. AJ is partially supported by the Department of Molecular Biosciences, Wenner-Gren Institute at Stockholm University. TMH was funded by The Wellcome Trust Centre for Cell Biology core funding (203149).

## Author contributions

Conceptualization: E.E. and N.V.; experiments: E.E., A.J., and S.J.; data analysis: E.E., M.T., J.P., I.S., A.J., and S.J.; software: T.M.H.; writing – first draft: E.E. and N.V.; writing – review and editing: E.E., M.T., J.P., I.S., S.J., A.J., and N.V.; funding acquisition: N.V. All authors read the manuscript.

## Declaration of interests

The authors declare no competing interests.

## Methods

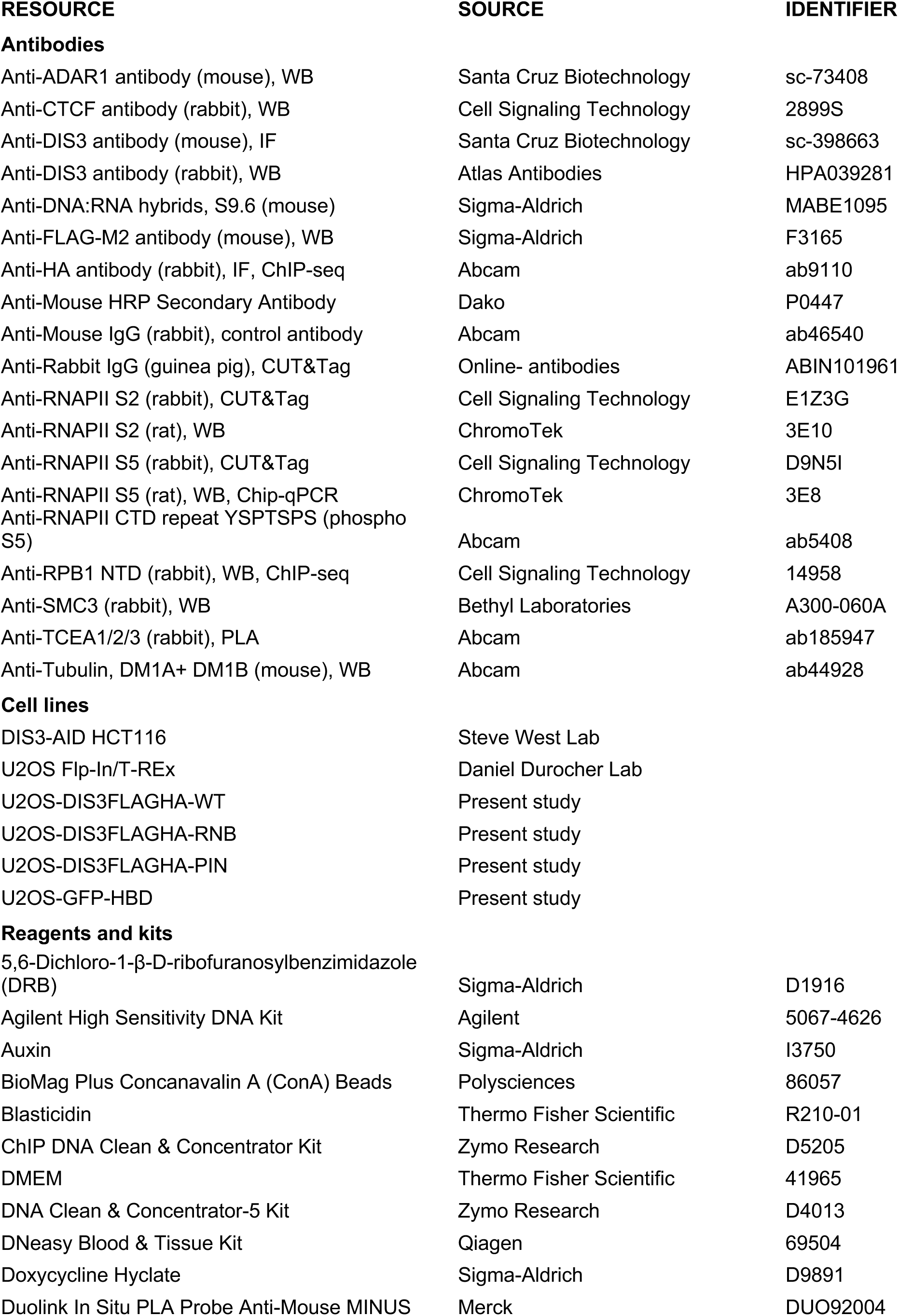
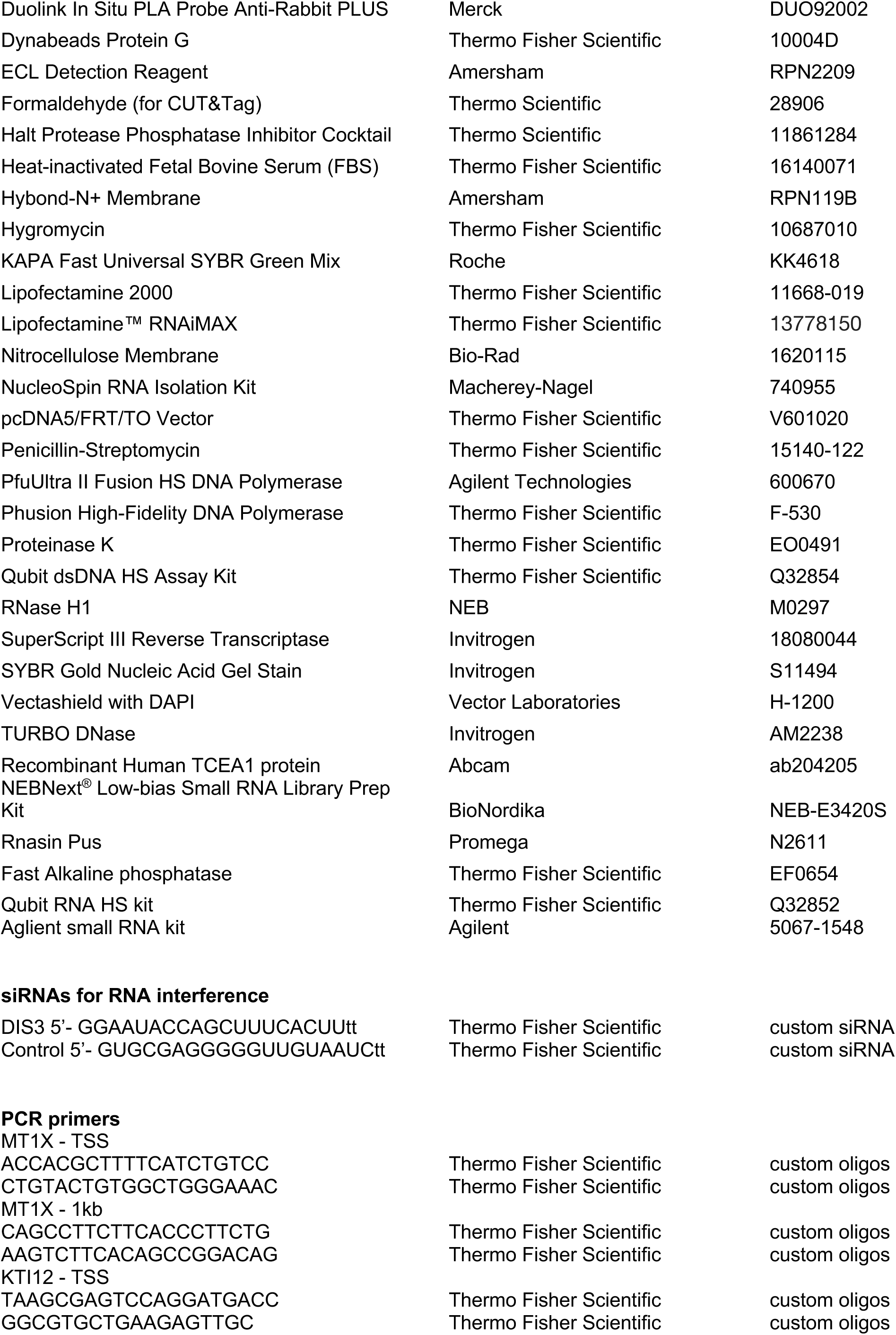
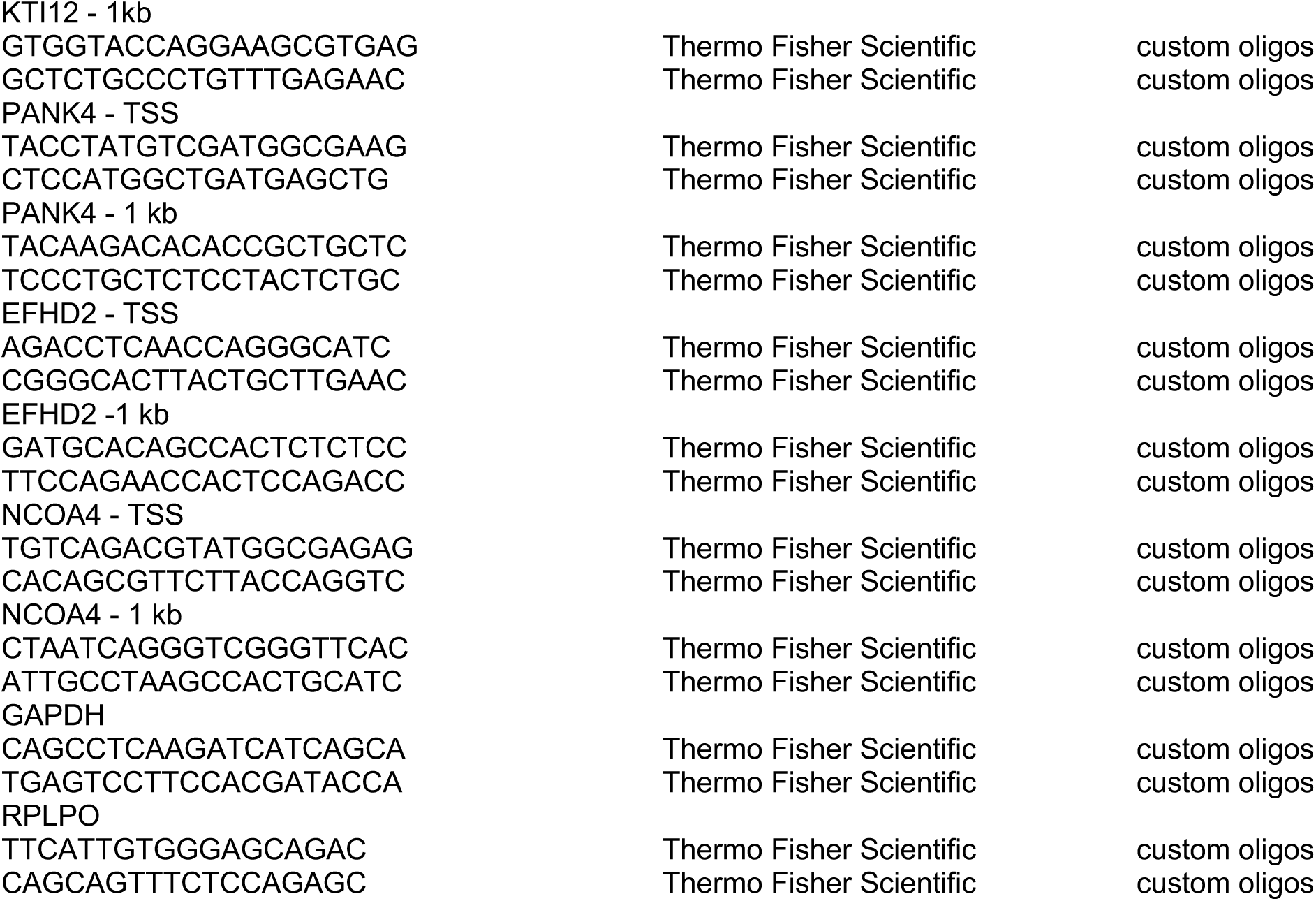

## Methods

### Cell lines and cell culture

U2OS and DIS3-AID HCT116 cell lines were cultured at 37°C in a humidified incubator with 5% CO2, in DMEM (#41965, Thermo Fisher Scientific) supplemented with Penicillin and Streptomycin (#15140-122 Thermo Fisher Scientific) and 10% of heat-inactivated fetal bovine serum (#16140071, Thermo Fisher Scientific). U2OS Flp-In/T-REx cells carrying doxycycline-inducible DIS3^FLAGHA^ variants were grown as above in the presence of 4 μg/ml Blasticidin (#R210-01, Thermo Fisher Scientific) and 100 μg/ml Hygromycin (#10687010, Thermo Fisher Scientific). All cell lines are routinely tested negative for mycoplasma. DIS3-AID HCT116 and U2OS Flp-In/T-Rex cell lines were kindly provided by Steven West (Univ of Exeter, UK) and Daniel Durocher (Univ. Toronto, Canada), respectively. For DIS3 depletion, DIS3-AID HCT116 cells were treated with 500 μM Auxin (#I3750, Sigma-Aldrich) or DMSO (carrier control) for 1 h unless otherwise indicated. In some cases, cells were irradiated with UV-C as indicated in th12 J/m^2^ or 20 J/m^2^ UVC as indicated.

### DIS3 depletion by RNA interference (RNAi)

Cells were transfected with siDIS3 or siCtrl using Lipofectamine™ RNAiMAX Transfection Reagent (Thermo Fisher Scientific). The cells were harvested and analyzed 48 h after transfection. Knock-down efficiencies were analyzed by Western blotting. SiRNA sequences are provided in the Resources list above.

### 5-Ethynyl uridine (5-EU) transcription assays

Steady-state transcription measurement in DIS3-AID cells was performed with cells treated with 500 μM Auxin (#I3750, Sigma-Aldrich) or DMSO for 1h, 4h or 20h, and labeled for 1h in the presence of 5-EU prior to fixation. For assessing nascent transcription in cells expressing exogenous DIS3^FLAGHA^ variants, the cells were grown in the presence of doxycycline for 48h before 5-EU labeling with 400 μM 5-EU in fresh medium for 20 min. The cells were fixed with 3.7% formaldehyde in PBS (FA; #252549, Sigma-Aldrich) for 15 min at RT and permeabilized with 0.5% Triton X-100 in PBS for 15 min at RT. Click-it-chemistry-based azide coupling was carried out using the Click-iT RNA Alexa Fluor 488 Imaging Kit (#C10329, Life Technologies) according to the manufacturer’s instructions with 1h incubation. To identify the cells expressing DIS3^FLAGHA^ variants, the Click-it chemistry was followed by immunofluorescent staining with anti-HA (ab9110, Abcam) or anti-DIS3 antibodies (sc-398663, Santa Cruz Biotechnology), as indicated in the figures. Coverslips were mounted using VectaShield with DAPI (#H-1200, Vector Laboratories). For the DRB wash-out assay, DIS3-AID cells were treated for 2h with 100 μM of 5,6-dichloro-1-b-D-ribofuranosylbenzimidazole (DRB, #D1916, Sigma-Aldrich) followed by incubation in the presence of either 500μM Auxin (#I3750, Sigma-Aldrich) or DMSO for 3h with Auxin or DMSO, still in the presence of DRB. The cells were washed and incubated for 20 min in fresh medium containing 5-EU for labelling of nascent RNA. For the DRB block assay, DIS3-AID cells were pre-treated for 3h with 500μM Auxin or DMSO followed by incubation with 100 μM DRB for 10 min and subsequent 5-EU labeling for 20min.

### Image analysis and quantification

Cells were imaged using a LSM700 (Zeiss) confocal microscope with ZEN 2011 black edition v7.0.5.288 software equipped with a ×40 Plan-Apochromat 1.3 NA oil-immersion lens or alternatively with a Plan-Apochromat 63x/1.40 oil DIC M27 objective. Images were analyzed and processed with Fiji/ImageJ2 v2.14.0/1.53o. DIS3^FLAGHA^ expression and nascent transcription (5-EU labelling) were quantified in individual cell nuclei with a manually set ROI inside the nucleus, based on DAPI staining, but avoiding the nucleolus. The 5^th^ and 95^th^ percentiles were included in the analysis. Relative fluorescence intensities were plotted using Prism/GraphPad software.

### CUT&Tag sequencing

CUT&Tag was based on Kaya-Okur et al. (2019) and Kaya-Okur et al. (2020) with some modifications. DIS3-AID cells were treated for Auxin or DMSO before irradiateion with 8 J/m^2^ UVC. Cells were washed in PBS, resuspended in ice-cold NE1 buffer (20mM HEPES-KOH pH 7.9, 10mM KCl, 0.5mM spermidine, 0.1% Triton X-100, 20% glycerol) supplemented with 1x Halt Protease and Phosphatase inhibitor cocktail (#11861284, Thermo Scientific), and incubated 10 min on ice. Nuclei were collected by centrifugation, resuspended in 0.1% formaldehyde (#28906, Thermo Scientific) in PBS and fixed for 2 min at RT. Fixation was quenched by adding 1.25 M glycine to a twice the molar concentration of formaldehyde. The cells were collected and resuspended in wash buffer (20mM HEPES-KOH pH 7.5, 150mM NaCl, 0.5M spermidine) to a concentration of 1x10^6^ cells /ml. BioMag Plus Concanavalin A (ConA) beads (#86057, Polysciences) were pre-washed in binding buffer (20mM HEPES-KOH pH 7.9, 10mM KCl, 1mM CaCl2, 1mM MnCl_2_) and 10 μl beads were added to each sample of 100.000 cells and mixed gently for 10 min at 4°. Nuclei-ConA bead complexes were collected and resuspended in 100 μl of antibody buffer (wash buffer containing 0.4mM EDTA and 0.1% BSA). 1.5 μl of primary antibody (S2 RNAPII E1Z3G or S5 RNAPII D9N5I) was added to each sample and incubated overnight at 4°C with gentle rocking. A no-primary-antibody sample was processed in parallel as negative control. Secondary antibody (guinea pig anti-rabbit antibody) diluted 1:100 was incubated with the nuclei-ConA bead complexes at RT for 1h. The rest of the CUT&Tag protocol was pwerfomred according to Kaya-Okur et al. (2019) with the exception of Tagmentation buffer used at a volume of 200 μl. The DNA was extracted using a DNA Clean & Concentrator-5 kit (#D4013, Zymo Research) and eluted in 21 μl of pre-warmed elution buffer. For the PCR and library enrichment, 20 μl of DNA eluate was used in a 60 μl reaction volume using 2x Phusion High-Fidelity PCR/GC Buffer master mix (#M0532S, NEB), with universal i5 primer and uniquely barcoded i7 primer. PCR cycles were as followed: Cycle 1: 58 °C for 5 min, Cycle 2: 72 °C for 5 min, Cycle 3: 98 °C for 30 sec, Cycle 4: 98 °C for 10 sec, Cycle 5: 63 °C for 10 sec Repeat Cycles 4-5 18 times. 72°C for 1 min and hold at 8°C. Post-PCR clean-up was performed using 0.8x volume of SPRI bead slurry per sample. After 10 min incubation at RT and two washes in 80% ethanol, elution was carried out in 15 μL of 10 mM Tris-HCl pH 8. Quality and quantity of libraries were assessed on 2100 Bioanalyzer (Agilent) using High Sensitivity DNA Kit (#5067-4626, Agilent), and on Qubit 2.0 using Qubit dsDNA HS Assay Kit (#Q32854, Thermo Fisher Scientific). The libraries were sequenced by the Bioinformatics and Expression Analysis core facility (BEA), Karolinska Insitute, on a Nextseq 2000 P2 100 Flowcell. Sequencing statistics are provided in Suppl. Table S1.

### Analysis of CUT&Tag data

CUT&TAG data was preprocessed using the nf-core cutandrun pipeline (v3.0) using the following parameters: nextflow run nf-core/cutandrun -r 3.0 SAMPLE_SHEET.csv --genome GRCh38 --use_control false --normalisation_mode CPM --peakcaller seacr,macs2 --consensus_peak_mode group --replicate_threshold 2 --minimum_alignement_q_score 20 --normalisation_binsize 1 --blacklist hg38-blacklist.v2.bed --validate_params= false. BAM files were converted to bigWig files using bamCoverage with parameters -b $FILE -o $FILE\.<u>scaled.bw</u> -p max --normalizeUsing CPM and all three replicates were combined using bigWigMerge. Subsequently, bigWig files were mapped to genes using deepTools as indicated: computeMatrix scale-regions -S BIGWIG_FILE -R BED_FILE --beforeRegionStartLength 1000 --regionBodyLength 3000 --afterRegionStartLength 1000 --skipZeros - o OUTPUT_FILE. These files were further processed in base R as indicated. For gene expression, preprocessed data from GEO accession GSE30786 was used. Gene lengths were taken from RefSeq. Elongation indices were calculated as log2 ratio of reads mapping to the terminal half of the gene body (adjacent to the TES) to the starting half of the gene body (adjacent to the TSS). P-values indicate results of a one-sided Mann-Whitney U test / Wilcoxon rank-sum test.

### Chromatin fractionation for Western blotting

Cells for chromatin fractionation was collected by trypsinization, pre or at indicated time points following UVC irradiation (20 J/m^2^) with 1h treatment with 500μM Auxin (#I3750, Sigma-Aldrich) or same volume of DMSO prior to UV. Chromatin-enriched fractions were prepared by incubating cells 5 min on ice in a cold hypotonic buffer, buffer A (10 mM HEPES, pH 7.9, 10 mM KCl, 1.5 mM MgCl2, 0.24M sucrose, 10% glycerol and 0.1% Triton X-100) supplemented with 1x Halt protease and phosphatase inhibitor (#1861281, Thermo Scientific). Following centrifugation for 4min 1300g at 4°C, the nuclei pellet was washed once in buffer A without detergent. The nuclei were lysed in buffer B buffer (3mM EDTA, 0.2 mM EGTA 1mM DTT) supplemented with 1x Halt protease and phosphatase inhibitor) and five rounds of snap-freezing cycles in liquid nitrogen and incubated on ice for 15min. Following centrifugation for 4min 1700g at 4°C, the pellet containing the insoluble nuclear extract and chromatin were washed once in buffer B and resuspended in Laemmli buffer, homogenised using a 29-gauge insulin syringe (#324891, VWR) and boiled for 10min.

### Western Blotting

Cells for Western blotting were collected, rinsed in PBS and lysed directly in Laemmli buffer. Samples were homogenised using a 29-gauge insulin syringe (#324891, VWR) and boiled for 10min. Proteins were separated by standard SDS-PAGE gel electrophoresis and transferred to nitrocellulose membrane (#1620115, Bio-Rad) overnight at 40V in transfer buffer adjusted for large proteins (390mM Glycine, 48mM Tris, 0.1% SDS and 20% methanol). Membranes were blocked in 5% milk in PBS containing 0.05% Tween-20 (PBST) and incubated with primary antibodies overnight at 4°C. Secondary antibodies were coupled to IRDyes (LiCor) and imaged with an Odyssey Fc infrared scanner (LiCor).

### Plasmid constructs

The coding region of the *DIS3* gene was amplified by PCR using Phusion High Fidelity DNA Polymerase (#F-530, Thermo Scientific) in four different segments using available internal restriction enzyme sites. Fragments were inserted sequentially into a pUC18 plasmid (#50004, addgene) modified to carry a compatible cloning cassette. Sequencing was performed after each insert to verify correct sequence. Site-directed mutagenesis (SdM) using PfuUltra II Fusion HS DNA polymerase (#600670, Agilient Technologies) was performed to generate resistance to knock-down by siRNA, an endoribonuclease deficient mutant (PIN mutant, D146N) and an exoribonuclease deficient mutant (RBD mutant, D487N), as well as a double D146N, D487N mutant. Full-length FLAG-HA C-terminally tagged Dis3 (DIS3^FLAGHA^) was transferred into pcDNA5/FRT/TO (#V601020, Thermo Fisher Scientific) using *Kpn1* and *XhoI* restriction enzymes. Primer sequences for Dis3 cloning and SdM are available upon request.

### Generation of doxycycline- inducible DIS3**^FLAGHA^** and GFP-HBD cell lines

To generate stable U2OS Flp-In cell lines carrying doxycycline-inducible DIS3^FLAGHA^- variants or doxycycline-inducible GFP-HBD, 600.000 U2OS Flp-In/T-REx cells were co-transfected with 4μg pcDNA5/FRT/TO-hDIS3^FLAGHA^ or pcDNA5/FRT/TO-GFP-HBD and 1μg the Recombinase-expressing plasmid pOGG44 using Lipofectamine 2000 (#11668-019, Thermo Fisher Scientific). At 48h post transfection, transfected cells were plated at low density and selected on 100μg/ml Hygromycin (#10687010, Thermo Fisher Scientific) and 4μg/ml Blasticidin (#R210-01, Thermo Fisher Scientific) for 9-12 days, after which individual clones were isolated, expanded and screened by immunofluorescence for doxycycline- induced expression of exogenous DIS3^FLAGHA^ or GFP-HBD.

### ChIP-sequencing of DIS3^FLAGHA^ and RPB1 NTD

U2OS Flp-In/T-REx cells harboring a doxycycline-inducible wild-type version of DIS3 was cultured in the presence of Doxycycline Hyclate (#D9891, Sigma) to a final concentration of 0.5μg/ml for 36h followed by a 12h culturing in the absence of doxycycline (36hON/12hOFF scheme). UV-irradiation was performed with a dosage of 8J/m^2^ and cells were collected pre-UV and at indicated timepoints following UV. Cells were cross-linked in 1% formaldehyde (#252549, Sigma-Aldrich) at 37°C for 20min, followed by 5 min quenching in 0.125 M glycine at RT. Immunoprecipitation was performed after pre-clearing with either α-HA (ab9110 abcam), α-RPB1 NTD (#14958, Cell Signaling) or IgG only control (ab46540, abcam). 10% of chromatin was isolated as input control. Dynabeads protein G slurry (#10004D, Thermo Fisher) were used to isolate the immunoprecipitated chromatin. Crosslinking reversal and DNA purification were carried out using standard procedures using a ChIP DNA Clean and Concentrator kit (#D5205, Zymo Research). Sample libraries were prepared by the National Genomics Infrastructure (NGI) at SciLifeLab using SMARTer ThruPLEX DNA-seq chemistry. Libraries were sequenced on NovaSeqXPlus (control-software 1.1.0.18335) with a 151nt(Read1)-19nt(Index1)-10nt(Index2)-151nt(Read2) setup using ’10B’ mode flowcell. The Bcl to FastQ conversion was performed using bcl2fastq_v2.20.0.422 from the CASAVA software suite. The quality scale used is Sanger / phred33 / Illumina 1.8+. Sequencing statistics are provided in Suppl. Table S2.

### Analysis of ChIP-seq data

ChIP-seq raw reads were pre-processed with nf-core/chipseq v1.2.2 (https://nf-co.re/chipseq/1.2.2) executed with Nextflow v21.10.6. Briefly, raw FASTQ files were quality-checked with FastQC (v0.11.9). Adapters were trimmed with trim-galore (v0.6.4) and reads were aligned to GRCh38 genome with Burrows-Wheeler alignment tool (BWA, v0.7.17) [https://doi.org/10.1093/bioinformatics/btp324]. PCR duplicates and multimapping reads were marked and removed with picard tool-suite (v2.23.1) and samtools respectively (v1.10). Reads overlapping blacklisted regions of the genome were discarded using bedtools (v2.29.2) and samtools. Normalised signal tracks scaled to 1 million mapped reads were generated with bedtools and bedGraphToBigWig function from ucsc-utilities. Output files such as bam and bigwig from nf-core pipeline were used for further analysis. A suite of Python tools known as deepTools (version 3.5.6) was used to generate metagene profiles from ChIP-seq data. BAM and BigWig files produced by the nf-core pipeline were used as input. The deepTools modules computeMatrix, plotProfile, and plotHeatmap were utilized to create the metagene profiles. Heatmaps were generated using deepTools (plotHeatmap), applying K-means hierarchical clustering.

### Analysis of backtracked RNA by LORAX-seq

LORAX-seq was performed according to Yang et al. (2024) with the following modifications. RNAPII complexes were immunoprecipitated with α-Rpb1 CTD (ab5408, Abcam) and captured using 1:1 mix of Dynabeads Protein-A and G. The magnetic beads were treated with recombinant Human TCEA1 protein (ab204205, Abcam) as per Yang et al. (2024), followed by isolation of eluted RNA using TRIZOL (Invitrogen). RNA was prepared for sequencing using the NEBNext Low-bias Small RNA library prep kit (NEB). The eluted RNA was mixed with QIAseq miRNA Library QC Spike-in (331535, Qiagen) as per manufacturer’s instructions. Sequencing libraries were visualized using an Agilent 2100 Tapestation and sequencing was done using the NextSeq 2000 P2 cartridge (Illumina). LORAX-seq raw reads were pre-processed with nf-core/rnaseq v3.21.0 (https://nf-co.re/rnaseq/3.21.0/) executed with Nextflow v25.04.7. Briefly, raw FASTQ files were quality-checked with FastQC (v0.12.1), adapters trimmed with trimgalore (v0.6.10), and reads were aligned to GRCh38.p14 using STAR (v2.7.11b). Alignment files were converted to bigwig format with spike-in normalization using samtools and deepTools (v3.5.5). PCR duplicates were marked using picard (v3.1.1), and read lengths were extracted using samtools (v1.20). Sequencing statistics are provided in Suppl. Table S3. The read length distribution plots were generated using R (v 4.4.2) for unique mapping reads. The deepTools (v3.5.5) modules computeMatrix were used to generate metagene profiles using normalized BigWigs files. Expressed genes were identified as described in CUT&Tag analysis (cutoff cutoff >10 fpkm). Read density was calculated by binning the normalized reads around TSS.

### RNA isolation, cDNA synthesis and qPCR

RNA was extracted from cells using using the NucleoSpin RNA isolation kit (#740955 Macherey-Nagel) following the manufacturer’s protocol. For reverse transcription, cDNA was synthesized using Random Hexamers and SuperScrip III reverse transcriptase (Invitrogen, #18080044). qPCR was carried out in a Mic qPCR cycler (Bio Molecular Systems) using 2× KAPA Fast Universal SYBR Green mix (#KK4618, Roche). Primer design was according to MIQE guidelines. All primer pairs fulfilled quality criteria regarding amplification efficiency and melting curves.

### DNA:RNA slot-blot assay

DIS3-AID cells were treated for 1h with either 500μM Auxin (#I3750, Sigma-Aldrich) or DMSO prior to exposure to 12J/m^2^ of UVC light. Samples were collected at indicated timepoints following UV irradiation and washed in cold PBS. gDNA was purified using DNeasy Blood and Tissue kit (# 69504, Qiagen) according to the manufacturer’s protocol with the following changes: the optional RNase A treatment step was omitted, cell lysis was performed at 56°C for 20min shaking and the final elution was done with 50+20μl elution buffer. Control samples were treated with RNase H1 (#M0297, NEB) for 2h at 37°C. 1μg of DNA per sample is blotted onto a wet Hybond-N+ membrane (#RPN119B, Amersham) using a slot-blot apparatus (Bio-Rad). Crosslinking to membrane was done using a UV Stratalinker 1800. The membranes were blocked with 5% milk/PBST for 30min and incubated with S9.6 antibody (#MABE1095, Sigma-Aldrich). An anti-mouse HRP was used as a secondary antibody (#P0447, Dako) and detection was performed using ECL detection reagent (#RPN2209, Amersham). Membranes were stained with SYBR Gold Nucleic Acid Gel Stain (S11494, Invitrogen) and imaged using a LiCor system.

### HBD-GFP retention assay by flow cytometry

HBD-GFP expression in U2OS Flp-In/T-REx cells was induced with 1 μg/ml Doxycyline Hyclate (#D9891, Sigma-Aldrich) for 48 h. The RNAi procedure was carried out as described. Cells were collected at indicated timepoints following UVC-light (20 J/m^2^) with trypsin and washed with 2% FBS/PBS followed by a pre-extraction with 0.05% Triton X-100 in PBS for 3 min at room temperature. Following a brief wash in 2%FBS/PBS cells were fixed with 4% paraformaldehyde for 10min at RT. Following a wash in 2%FBS/PBS, cells were analyzed on an FACSMelody (Becton Dickinson) where 100,000 cells were measured per sample. Data analysis was performed with FlowJo (v10.10.01, Becton Dickinson). After gating for SSC singlets (SSC-W/SSC-H), FSC singlets FSC-W/FSC-H) and DAPI-A/Hoechst-A), the proportion of GFP- positive cells was quantified by setting a threshold above the fluorescence of No Dox control cells.

### Proximity ligation assay (PLA)

DIS3-AID cells were fixed in 4% paraformaldehyde in PBS, pH 7.4 and permeabilized with 0.5% Triton X-100 in PBS for 20 min at room temperature. The antibodies used were anti-DIS3 (Santa Cruz Biotechnology, sc-398663) and anti-TCEA1,2,3 (Abcam, ab185947). PLA was performed according to the manufacturer’s protocol (Duolink, Sigma Aldrich) using Duolink In Situ PLA Probe Anti-Mouse MINUS and Anti-Rabbit PLUS. The preparations were mounted using VectaShield with DAPI (#H-1200, Vector Laboratories). Negative control PLA reactions were carried out in parallel with only one primary antibody. Images were acquired using a LSM 800 AiryScan (G201K) laser confocal microscope (Zeiss) with a 63x oil immersion objective. Z-stacks were acquired consisting of 10 sections, 0.6 µm/section. The number of PLA foci/nucleus were counted in z-stack projections using the plugin Elin_Intensities_v2_1 (https://doi.org/10.5281/zenodo.17640956) in randomly selected areas of the preparations. The number of cells analyzed is indicated in the figure.

### Statistical testing

In bar plots, the bars show average values and the error bars represent standard deviations unless otherwise indicated. The number of biological replicates for each experiment and the statistical tests used in each case are indicated in the figure legends. Probability values (p_adj_) for statistically significance are provided in the figures.

## Supplementary material

Supplementary Figures S1-S7

Supplementary Tables S1-S3

## Notes

### Competing Interest Statement

The authors have declared no competing interest.

