## Supplemental Figures S1-S7 for "RNA degradation by DIS3 is a necessary step in the resolution of backtracked transcription complexes"

**Supplemental Figure S1. Overexpression of DOX-inducible DIS3 rescues the transcriptional defect induced by DIS3 depletion**

**Supplemental Figure S2. Effect of DIS3 depletion on RNAPII in different cell lines**

**Supplemental Figure S3. Analysis of DNA:RNA hybrids in DIS3-depleted cells**

**Supplemental Figure S4. ChIP-seq analysis showing RPB1 NTD and DIS3 distributions in clusters 1-3**

**Supplemental Figure S5. LORAX-seq analysis of backtracked RNAs in DIS3-AID cells**

**Supplemental Figure S6. Proximity ligation analysis of DIS3 and TFIIS in DIS3-AID cells**

**Supplemental Figure S7. Expression of DOX-inducible DIS3 variants in U2OS-DIS3$^{FLAGHA}$ cells**

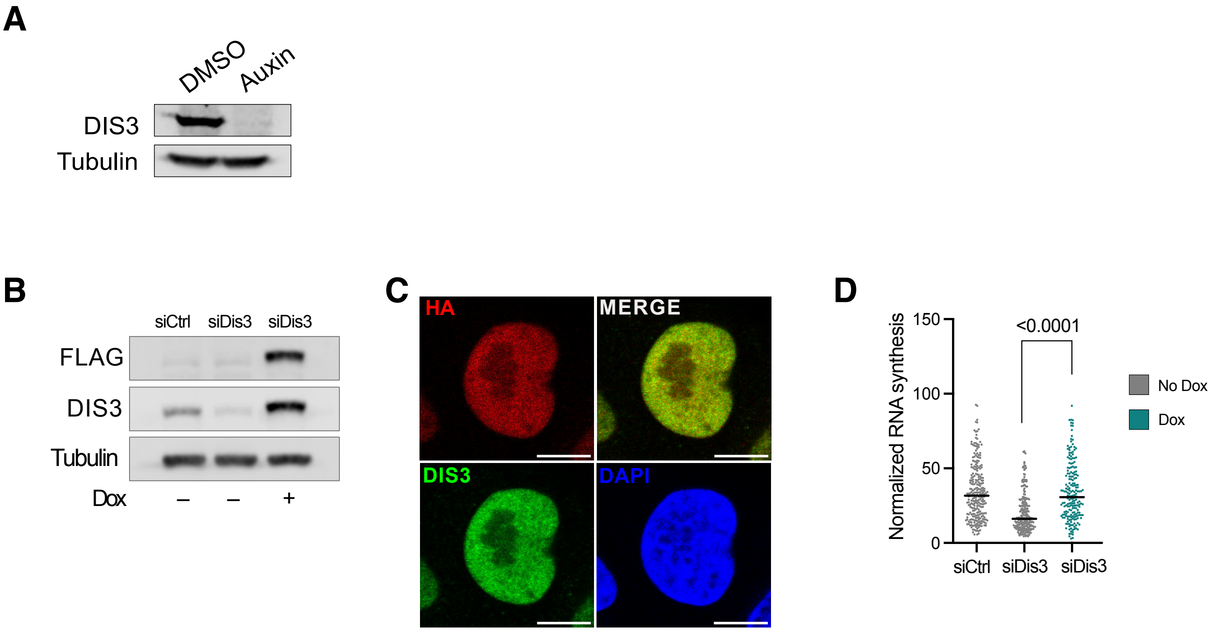


**Supplemental Figure S1. Overexpression of DOX-inducible DIS3 rescues the transcriptional defect induced by DIS3 depletion**

(A) Representative Western blot showing rapid depletion of DIS3 in DIS3-AID HCT116 cells. The cells were treated with either Auxin or DMSO (control) for 1 h. Tubulin was used as a loading control.

(B) Representative Western blot of U2OS FLP-In T-Rex cells harboring a siRNA-resistant wild-type version of DIS3 (U2OS-DIS3$^{FLAGHA}$-WT). U2OS-DIS3$^{FLAGHA}$-WT expression was induce by culturing the cells in the presence of 0.5 µg/ml doxycycline for 20 h. DIS3 was depleted by RNA interference using either siDis3 or control siRNA (siCtrl), as indicated. Tubulin was used as a loading control.

(C) Representative immunofluorescence images of a U2OS-DIS3$^{FLAGHA}$-WT cell expressing DIS3 WT. The scale bars represent approximately 10 µm.

(D) Analysis of nascent RNA by 5-EU incorporation in the indicated siCtrl and siDis3 knockdown cells. When indicated, cells were grown in the presence 0.5 µg/ml doxycycline for 48 h. The absolute intensity of the 5-EU labeling was normalized within each experiment. Black lines indicate the median. The figure includes data from 5$^{th}$ and 95$^{th}$percentile of all cells from three individual experiments. Number of cells analyzed in each condition: siCtrl -DOX: Pre *n*=263, 3h UV *n*=219, 20h UV *n*=262; siDis3 -DOX: Pre *n*=235, 3h UV *n*=248, 20h UV *n*=260; siDis3 +DOX: Pre *n*=200, 3h UV *n*=199, 20h UV *n*=270. A Mann-Whitney test was used to test statistical significance.

**A**

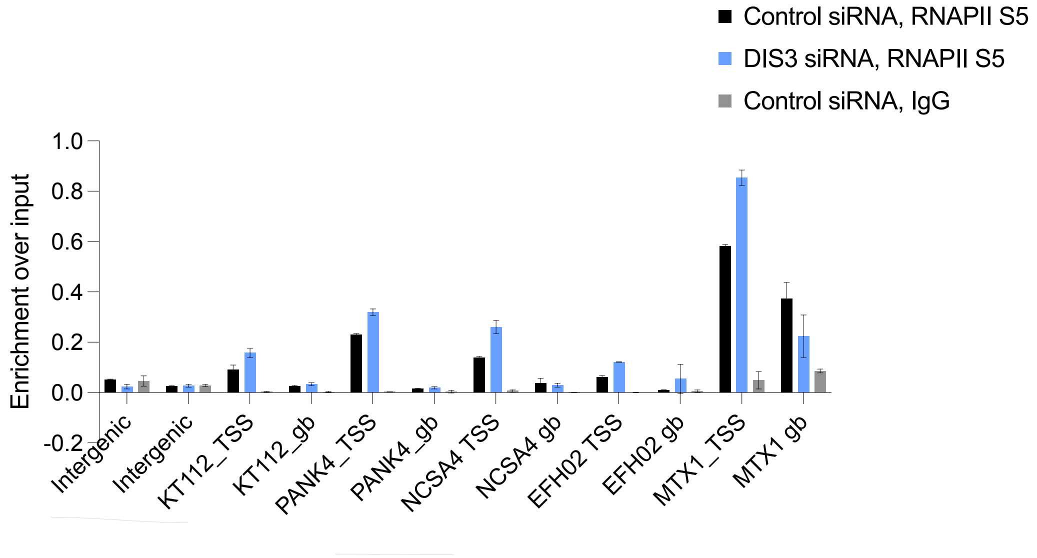


**B**

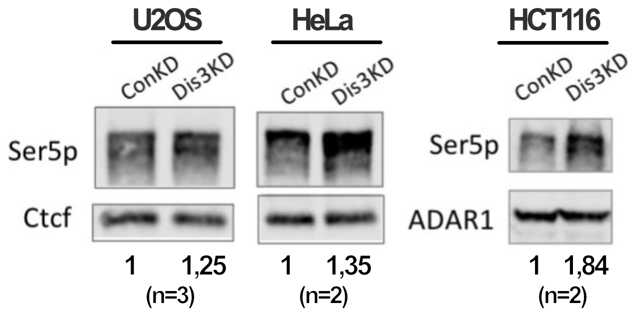


**Supplemental Figure S2. Effect of DIS3 depletion on RNAPII in different cell lines**

(A) U2OS cells were transfected with siRNAs (siDIS3 or siCtrl) to deplete DIS3, and the effect of the depletion on RNAPII occupancy was analyzed by ChIP-qPCR using the S5 RNAPII (3E8) antibody. Immunoprecipitated DNA was isolated and analyzed by qPCR using primers for selected genes, as indicated in the figure. For each gene, RNAPII S5 signal was analyzed at the TSS and 1 kb downstream (gb, gene body). Two intergenic regions were analyzed in parallel as negative controls to assess the specificity of the ChIP-qPCR signals. The bars show average enrichment relative to input from two technical replicates. The error bars represent standard deviations.

(B) Western blot analysis of chromatin-bound proteins extracted from different cell lines, as indicated, in control cells and DIS3-depleted cells. CTCF and ADAR1 were used as loading controls for normalization purposes. The figures under the blots show average enrichment in siDis3 relative to controls (siCtrl). The number of independent replicates is shown in the figure.

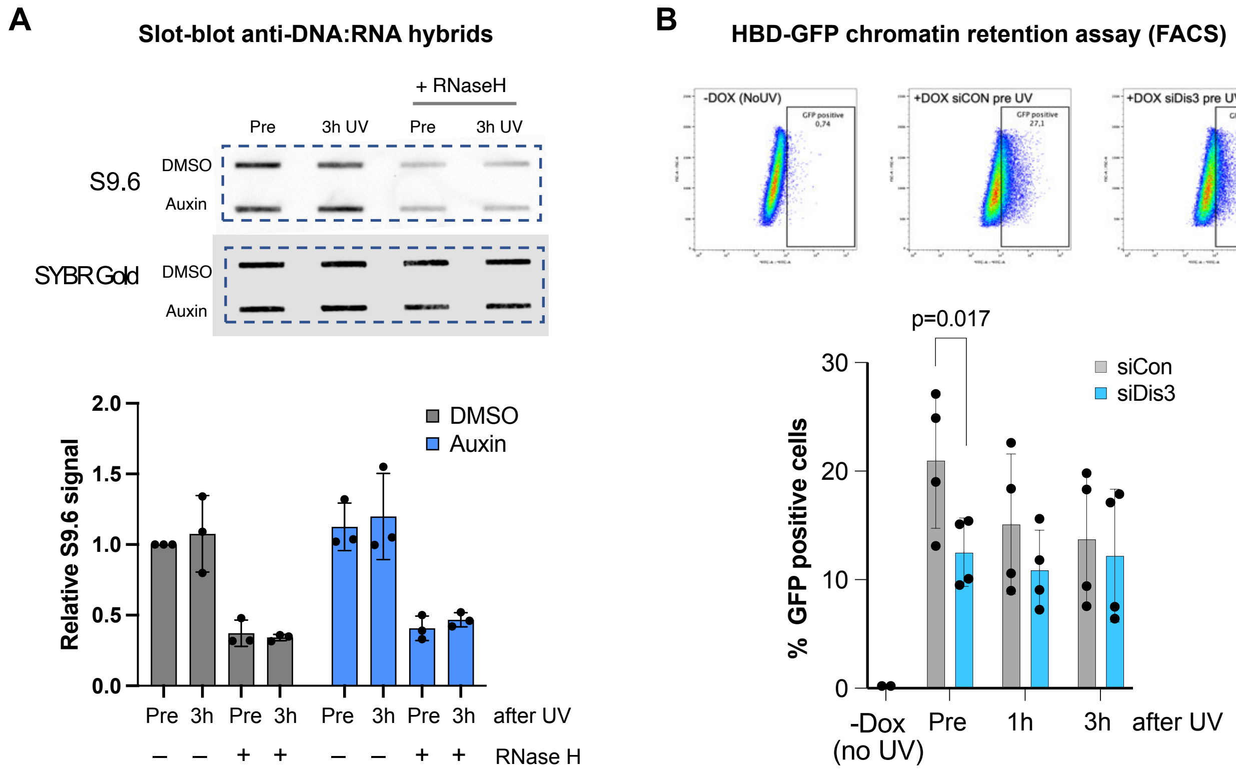


**Supplemental Figure S3. Analysis of DNA:RNA hybrids in DIS3-depleted cells**

(A) Slot blot analysis to quantify DNA:RNA hybrids in the genome of DIS3-AID cells using the S9.6 antibody. Cells were treated with either DMSO or Auxin for 1 h and irradiated with 12 J/m2 UVC. RNaseH was used to asses the specificity of the signals. The blots were stained with SYBR Gold to quantify total DNA for normalization purposes. The bars show relative fold changes in S9.6 signal compared to DMSO pre-irradiated samples. Error bars display standard deviations from three independent experiments. No significant changes in S9.6 signal were detected as a result of DIS3 depletion.

(B) Flow cytometry analysis to quantify DNA:RNA hybrids in the genome of U2OS Flp-In cells expressing a doxocycline-inducible RNase H binding domain fused to GFP (HBD-GFP). DIS3 was depleted by RNAi for 48h. In some cases, cells were irradiated with with 12 J/m2 UVC, as indicated. The binding of HBD-GFP to chromatin was analyzed by flow cytometry and the fraction of GFP-positive cells was used as a proxy for DNA:RNA hybrid levels. In control cells (pre UV), DIS3 depletion resulted in decreased HBD-GFP signal. Student t-test was used for statistical testing, n=4 independent experiments, 100 000 analyzed cells in each condition.

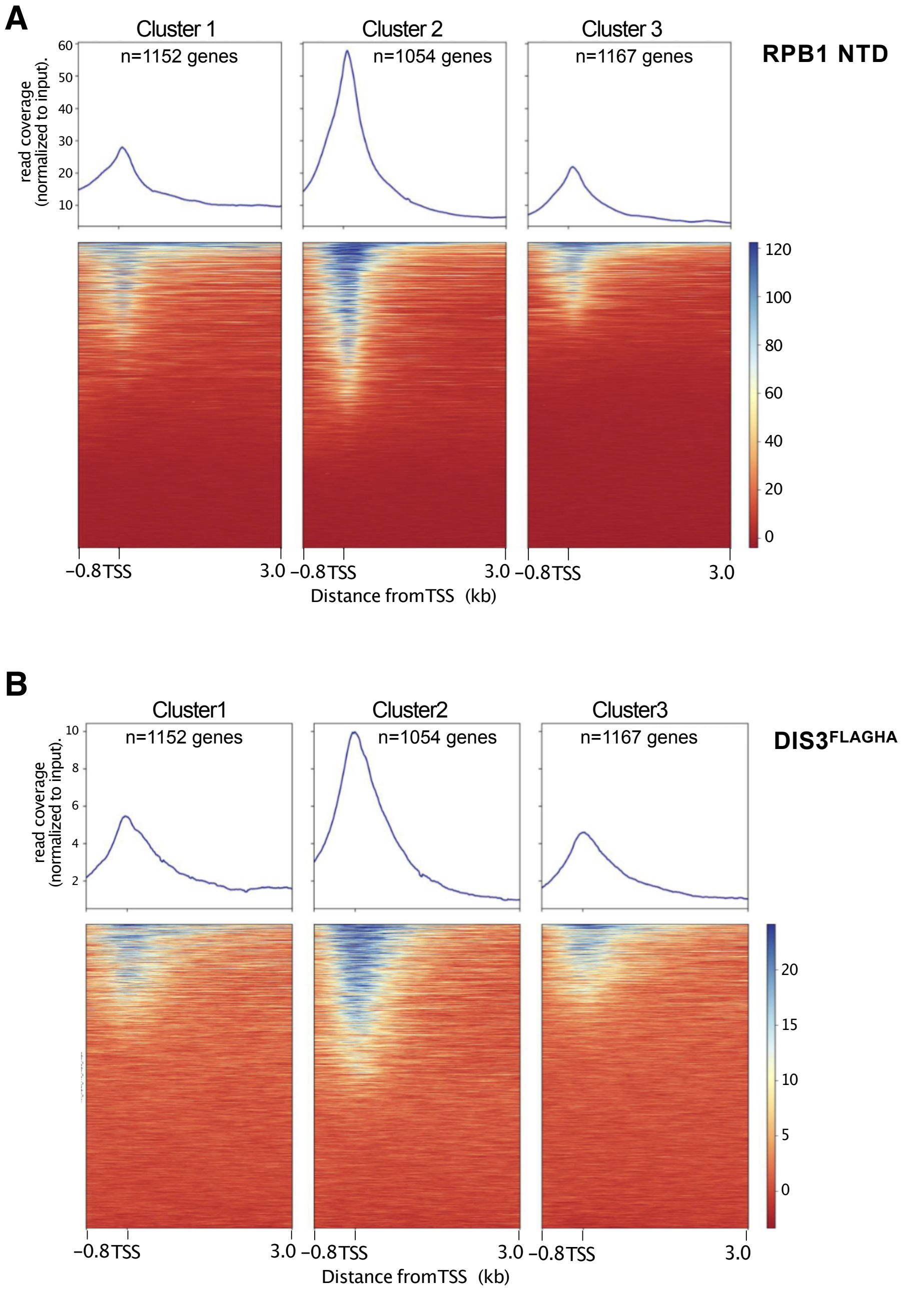


**Supplemental Figure S4. Chip-seq analysis showing RPB1 NTD and DIS3 distributions in clusters 1-3**

(A) Metagene plots and corresponding heat maps showing the distributions of RPB1 NTD at the TSS and 3 kb into the gene body.

(B) Metagene plots and corresponding heat maps showing the distributions of DIS3$^{FLAGHA}$-WT at the TSS and 3 kb into the gene body.

The metagenes show average read coverage normalized to input. The clusters are those defined in Fig.3C based on the RPB1 NTD change at 3 h after UV compared to pre UV. The plots include merged data from two replicates.

Cluster 2 shows the highest RPB1 and DIS3 levels.

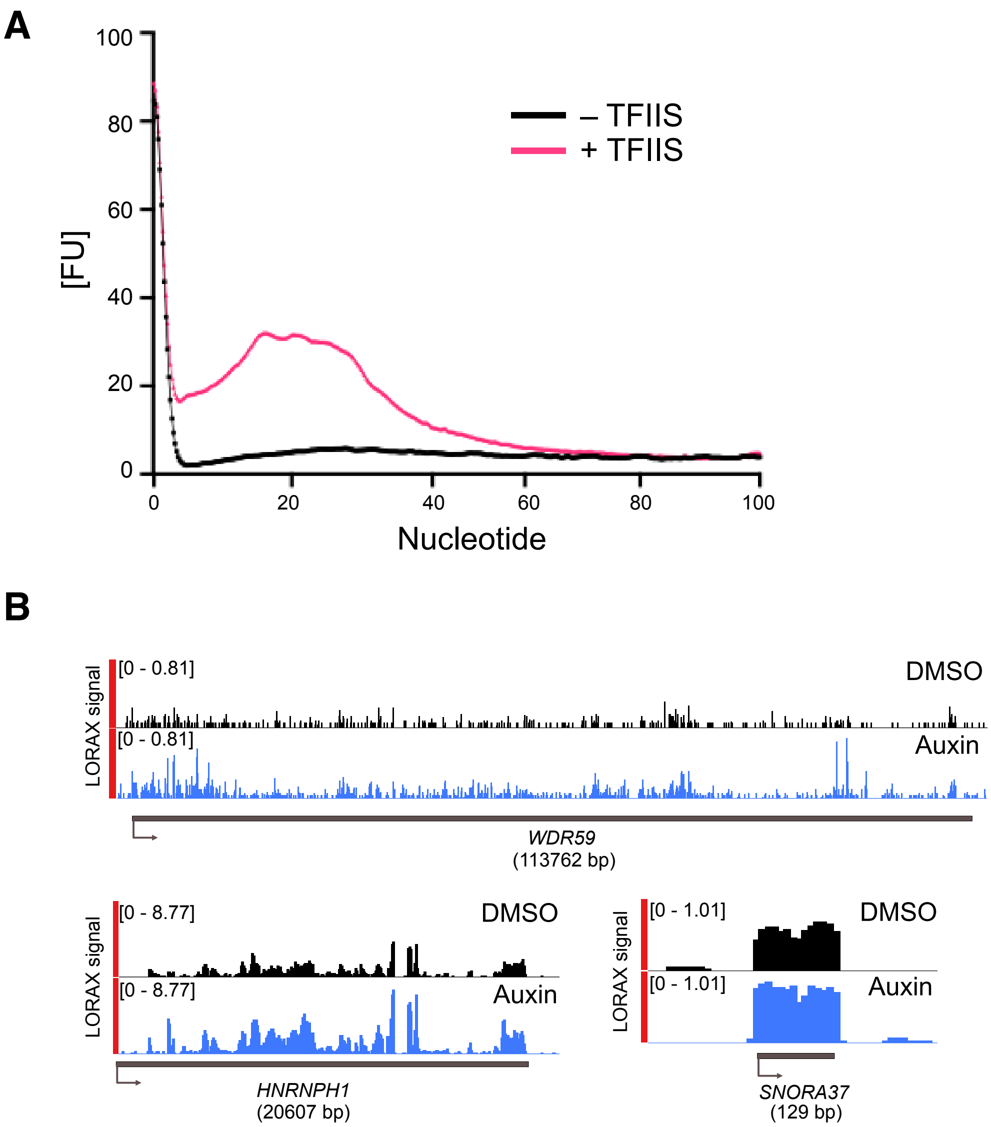


**Supplemental Figure S5. LORAX-seq analysis of backtracked RNAs in DIS3-AID cells**

A. The line plot depicts length distribution of cleavage products from LORAX-seq libraries prepared from DMSO (control) cells with and without TFIIS cleavage *in vitro*. The release of backtracked RNA products is seen in the +TFIIS samples. The plot shows average distributions from three independent replicates.

B. Representative genome viewer of normalized LORAX-seq signal across two protein-coding genes (*WDR59* and *HNRNPH1*) and a non-coding gene (*SNORA37*) in control cells (DMSO) and DIS3-depleted cells (Auxin). Gene lengths are indicated below gene names. The figure shows merged data from two replicates for each condition.

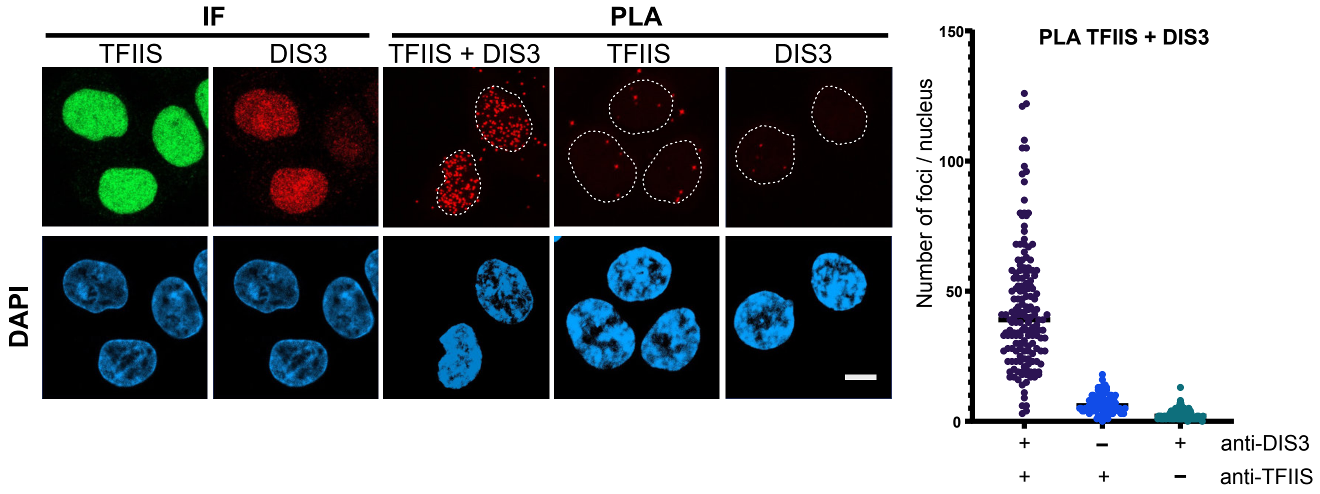


**Supplemental Figure S6. Proximity ligation analysis of DIS3 and TFIIS in DIS3-AID cells**

Immunofluorescence (IF) and proximity ligation assays (PLA) were performed in DIS3-AID cells using anti-TFIIS and anti-DIS3 antibodies. From left to right: IF images for TFIIS (green, upper panel) and DIS3 (red, upper panel); a representative PLA image showing TFIIS-DIS3 interaction; two negative-control PLA reactions performed with only one primary antibody (TFIIS only or DIS3 only; upper panels). The figure also shows the corresponding DAPI images (blue, lower panels). The scale bar represents approx. 10 µm. The plot on the right-hand side shows number of foci/nucleus in each condition, as indicated, from two independent experiments. Number of cells analyzed in each condition: n=165 (PLA TFIIS & DIS3), n=99 each (TFIIS only and DIS3 only).

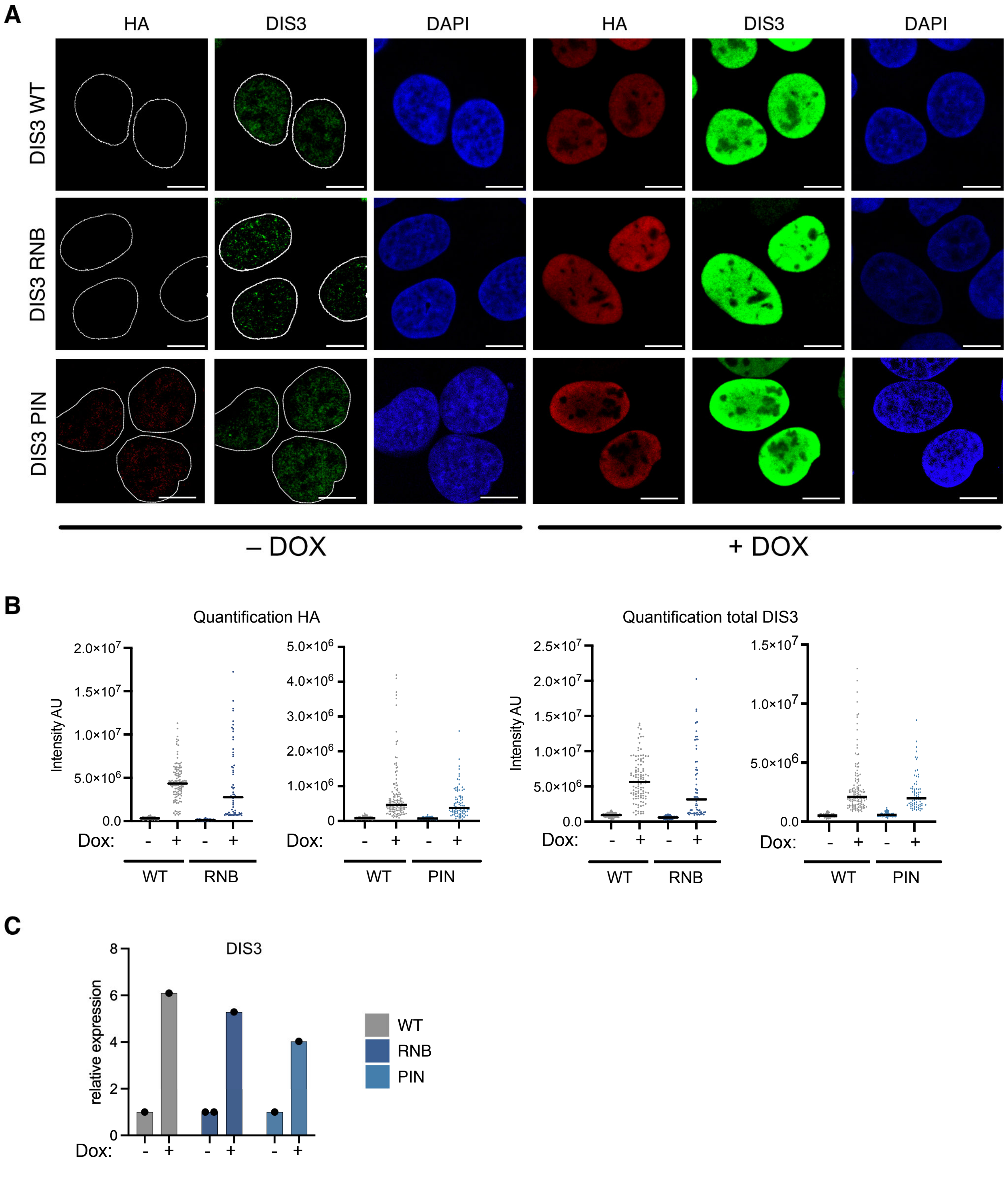


**Supplementary Figure S7. Expression levels of DOX-inducible DIS3 variants in U2OS-DIS3$^{FLAGHA}$ cells**

(A) Representative immunofluorescence images of U2OS FLP-In T-Rex cells expressing DIS3$^{FLAGHA}$ variants (DIS3$^{FLAGHA}$-WT, DIS3$^{FLAGHA}$-RNB and DIS3$^{FLAGHA}$-PIN RNB), as indicated, in the presence or absence of 0.5 µg/ml doxycycline for 48h. The scale bars represent 10 µm.

(B) Quantification of exogenous DIS3$^{FLAGHA}$ proteins (HA, left) and total DIS3 levels (DIS3, right) from (A). One representative experiment is shown out of three performed. To remove negative cells that failed to express the exogenous DIS3$^{FLAGHA}$ variant, an arbitrary threshold for HA positive (+Dox) and negative (-Dox) cells was set in each experiment. Only cells above (+Dox) or below (-Dox) the threshold were included in the analysis.

(C) Relative level of expression of the DIS3$^{FLAGHA}$ variants compared to the levels of DIS3 in -Dox cells for each individual cell line.
